# Tetherin enforces an immunometabolic checkpoint that coordinates glycolytic and interferon signaling in adipocytes

**DOI:** 10.64898/2026.07.06.735062

**Authors:** Chung Hwan Cho, YoungUk Jang, Aidan Warnock, Ramazan Yildiz, Jing Jhang, Kajal Davi, Niki F. Brisnovali, Victoria Huhn, Peng Wang, Romina Bevaqua, Leigh Goedeke, Michael A. Schotsaert, Mirela Berisa, Daniel Puleston, Prashant Rajbhandari

## Abstract

Coordination between innate immune signaling and glucose metabolism is fundamental to organismal homeostasis, yet despite decades of study linking immunity and metabolism, the mechanisms by which metabolic cells restrain antiviral innate signaling while preserving glycolytic competence during overnutrition remain poorly defined. Here we identify Tetherin (BST2) as a unique cell-intrinsic immunometabolic checkpoint that couples restraint of type I interferon (IFN-I) signaling to preservation of glycolytic capacity in adipocytes. Tetherin localizes to endoplasmic reticulum and organizes an interactome enriched for antiviral sensing regulators and glycolytic control nodes in adipocytes. Mechanistically, Tetherin directly engages the ubiquitin-dependent degradation machinery NDFIP1 and RNF128 to terminate IRF3 activation, thereby limiting pro-inflammatory, anti-glycolytic signaling and protecting adipocytes from metabolic dysfunction. In parallel, multiomics integration reveals that Tetherin also acts as a scaffold that binds and spatially organizes and activates PFKFB3 to increase glycolytic capacity and restrain MAVS–IRF3 innate immune signalling. In vivo, adipocyte-specific loss of Tetherin amplifies high sucrose diet and high-fat-diet–induced glucose intolerance and liver steatosis, whereas overexpression of human Tetherin in adipocyte suppresses obesity-driven interferon signaling, restores glycolytic pathway, and improves metabolic homeostasis. Orthogonal perturbations in cancer and insulinoma cells further confirm an immunometabolic role for Tetherin. Together, these findings define Tetherin as a dual node immunometabolic checkpoint that couples restraint of antiviral innate inflammatory signaling to maintenance of glycolytic competence, thereby safeguarding adipocyte metabolic homeostasis.

## Introduction

Obesity and diabetes are marked by a progressive failure of metabolic tissues to maintain homeostasis in the presence of nutrient excess, a process tightly linked to chronic low-grade inflammation, nutrient-sensing stress pathways and impaired insulin action ^1–3^. In adipose tissue, this failure is accompanied by inflammatory signalling, impaired glucose handling, reduced metabolic capacity, and secondary pathology in distal organs such as liver, including hepatic insulin resistance and steatosis ^3–7^. Although inflammatory pathways are widely implicated in these disorders, broad immunosuppression is neither mechanistically precise nor clinically ideal ^8^, highlighting the need to define endogenous tissue/cell-intrinsic mechanisms that restrain inflammatory signalling while preserving core metabolic function.

Adipocytes are now recognized as active immunometabolic cells that integrate nutrient sensing, endocrine signalling and inflammatory stress responses ^9–14^. Among the pathways engaged by cellular stress, the MAVS (Mitochondrial Antiviral Signaling Protein)–TBK1 (TANK-binding Kinase 1)–IRF3 (Interferon Regulatory Factor 3) axis is classically associated with antiviral defence and type I interferon production^15^. Yet chronic or unresolved activation of IRF3 and innate immune pathways in metabolic tissues can become maladaptive ^14,16–19^. In obesity, TBK1 and IKKε link inflammatory signalling to altered energy balance and insulin resistance ^19–22^, and mitochondrial stress–activated cGAS–STING signalling has been implicated in adipose inflammation, impaired thermogenesis and metabolic dysfunction ^23–25^. Type I interferon signalling also remodels core metabolism, including thermogenesis, fatty-acid oxidation and oxidative phosphorylation ^10,26–34^.

In adipocytes, how innate antiviral signalling is resolved, and whether its resolution is mechanistically coupled to maintenance of glycolytic competence remains unknown. This problem is important because glucose metabolism is a fundamental determinant of adipocyte function ^35^. Beyond ATP production, glucose flux supports insulin-stimulated glucose disposal, lipid synthesis, adipocyte differentiation, thermogenic capacity, and systemic metabolic homeostasis. Hexokinase 2 (HK2) commits glucose to intracellular metabolism, and diet-induced loss of adipose HK2 reduces adipocyte glucose disposal and contributes to hyperglycemia insulin resistance^36^. PFKFB3, a key regulator of fructose-2,6-bisphosphate and glycolytic flux, protects against diet-induced adipose inflammation and systemic insulin resistance^37^.

Here we show that Tetherin, also known as BST2, an interferon-inducible membrane protein originally defined as an antiviral restriction factor that prevents release of enveloped virions^38^, acts as an adipocyte-intrinsic immunometabolic checkpoint. BST2 suppresses tonic IRF3 activation and interferon target-gene induction while preserving glycolytic signalling and metabolite flux. Mechanistically, BST2 associates with degradation machinery, NDFIP1 and RNF128 to restrain innate immune signalling and interacts with PFKFB3 to promote a cytoplasmic, phosphorylated PFKFB3 state that supports glycolysis. In vivo, adipocyte BST2 improves glucose tolerance, insulin sensitivity, energy expenditure and protection from hepatic steatosis under dietary stress. Together, these findings identify a direct molecular coupling between immune-signal resolution and glycolytic competence in adipocytes and position BST2 as a regulator of systemic metabolic health.

## Results

### BST2 controls adipocyte glycolytic program

Obesity is associated with chronic activation of innate immune pathways in adipose tissues, including induction of type I interferon signaling and a broad interferon-stimulated gene (ISG) program. Transcriptomic mining of IFN-treated primary adipocytes ^33^ and obese human and murine single cell adipose atlases^39^ identified a small four-gene core of highly induced interferon-β–induced genes (ISGs) that is paradoxically suppressed in obese adipocyte and preadipocyte lineages: BST2, GREB1L, PTX3 and SOCS1 (Fig. S1A). Among these, BST2 (also known as Tetherin/CD317) was the most robustly and selectively suppressed in obese versus lean human adipocyte and preadipocyte fractions in the Emont et al. atlas ^39^ (adipocyte log₂FC = −0.85, pₐdⱼ = 2.2e^−02^; preadipocyte log₂FC = −0.97, pₐdⱼ = 2.8e⁻^04^; Fig. S1B). In high-fat-diet (HFD)-fed mice white adipose tissue (WAT), *Bst2* was ranked among the most suppressed ISGs in macrophages (log₂FC = −3.50, rank 2/1,399), preadipocytes (log₂FC = −1.44, rank 33/1,399) and monocytes (log₂FC = −2.36, rank 4/1,399) (Fig. S1C). Spearman correlation in human adipose tissues ^40^ linked BST2 expression negatively to BMI, HOMA-IR and waist circumference (Fig. S1E, left), and a differentiation time-course showed that BST2 protein rises sharply during D0–D6 of adipogenesis in MAPC- and TERT-immortalized preadipocyte lines^41^ (Fig. S1E, right; Fig. S1D, in-cell western validation across WT, BST2KO and BST2-overexpressing (BST2OE) differentiated 10T1/2 adipocytes).

BST2 is a type II transmembrane protein consisting of a short N-terminal domain followed by an alpha-helical transmembrane domain, a labile coiled-coil ectodomain, and a C-terminal glycosyl-phosphatidylinositol (GPI) anchor ^42^ that inhibits the release of budding virions from infected cells ^43^. BST2 was recently shown to mark certain pool of beige progenitors ^44^ and was also identified in human visceral WAT, where it correlated with a thermogenic signature ^45^. Beside the role of BST2 in virology, the function of BST2 protein in adipocyte, adipocyte inflammation, adipose tissue biology, and metabolism, is not known.

BST2 protein in adipocytes was present predominantly in the endoplasmic reticulum (ER): dual immunofluorescence with ER-Tracker showed punctate BST2 signal that co-localized with the ER network only in differentiated adipocytes and not in undifferentiated preadipocytes and, with the expected loss of BST2 signal in BSTKO adipocytes and amplified signal in BST2OE cells (Fig. 1A). In inguinal white adipose tissue (iWAT), BST2 was present at a basal level and was strongly induced by the β3-adrenergic agonist CL-316,243 and by cold exposure, where it concentrated at the periphery of Plin1⁺ lipid droplets in mature adipocytes (Fig. 1B). Fractionation of iWAT and gonadal WAT (gWAT) into stromal-vascular (SVF) and mature-adipocyte compartments confirmed that adrenergic stimulation enriched BST2 protein in the mature-adipocyte fraction (Fig. S1F,G). Together, these data identify BST2 as an adipocyte ER-localized ISG whose expression is increased under adrenergic conditions and decreased in obese adipose tissue, suggesting a metabolic role in adipose tissue.

**Figure 1.**
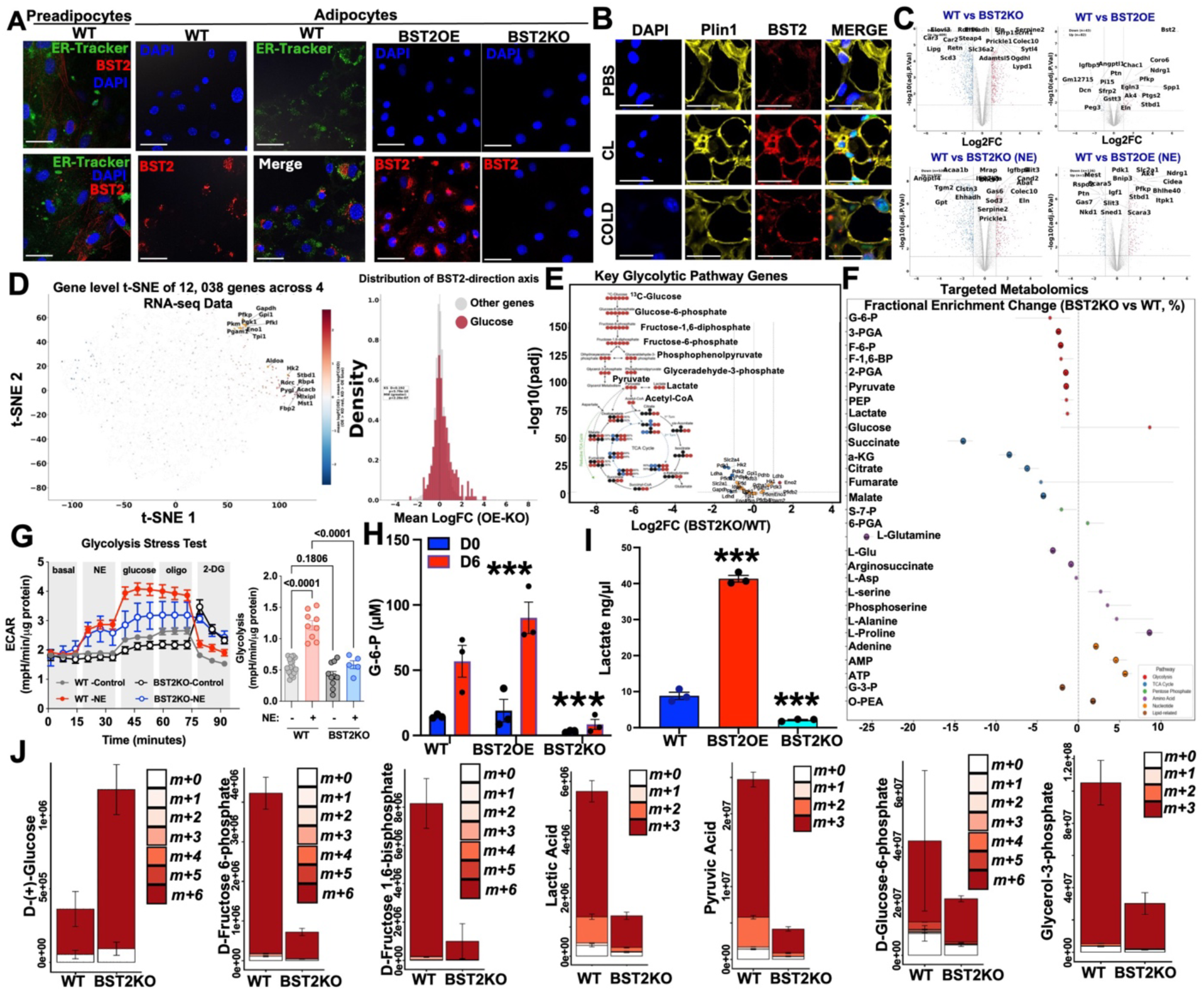
BST2 is an adipocyte ER-localized, adrenergically induced positive regulator of adipocyte glycolytic flux. (A) Immunofluorescence of WT 10T/1 preadipocytes and WT, BST2OE and BST2KO mature 10T1/2 adipocytes (adipocytes) co-stained for BST2 (red), ER-Tracker (green) and DAPI (blue). Punctate BST2 signal co-localizes with the ER network in WT and BST2OE cells and is absent in BST2KO. Scale bar, 50 µm. (B) IF of iWAT sections stained for DAPI, Plin1 (lipid-droplet outline, yellow), BST2 (red) and MERGE under PBS (vehicle), CL-316,243 (β3-adrenergic agonist, 1 mg kg⁻¹ i.p., 48h) or 24 h cold exposure (6 °C). Scale bar, 100 µm. (C) Volcano plots (−log₁₀ pₐdⱼ vs log₂FC) for four bulk RNA-seq conditions: WT vs BST2KO and WT vs BST2OE, each at baseline and under NE (1 µM, 6 h). Genes with |log₂FC| > 1 and pₐdⱼ < 0.05 are colored (blue, down; red, up); representative metabolic gene labels are shown. (D) Gene-level t-SNE embedding of 12,038 expressed genes across the four RNA-seq conditions, colored by mean log₂FC(BST2OE − BST2KO). The marginal density of glucose-metabolism genes (red) is right-shifted relative to all other genes (grey): Kolmogorov–Smirnov D = 0.192, p = 5.79 × 10⁻⁷⁰; Mann–Whitney p = 2.26 × 10⁻⁷⁷. (E) Volcano plot of downregulated glycolytic genes from RNA-seq data in BST2KO versus WT day-6 adipocytes (−log₁₀ pₐdⱼ vs log₂FC(KO/WT)). Model of ¹³C-glucose-trace-relevant intermediates are annotated; key glycolytic species (glucose, G-6-P, F-6-P, F-1,6-BP, GAP, PEP, lactate, pyruvate, acetyl-CoA) are labelled. (F) Fractional-enrichment change (BST2KO vs WT, %) across ∼30 metabolites grouped by pathway (glycolysis, TCA cycle, pentose-phosphate pathway, amino acids, nucleotides, lipid-related). Bars are mean ± s.e.m. (n = 4 biological replicates per genotype). (G) Real-time Seahorse extracellular-acidification-rate (ECAR; mpH min⁻¹) glycolysis-stress test in Ctrl, Ctrl-NE, BST2KO and BST2KO-NE adipocytes. Sequential injections of NE, glucose, oligomycin and 2-DG are indicated. Data are mean ± s.e.m., n = 5–6 wells per condition. (H) Steady-state G-6-P concentration (µM) in WT, BST2OE and BST2KO adipocytes at D0 and D6 of differentiation. Bars are mean ± s.e.m. with individual replicates overlaid; ***p < 0.001 (two-way ANOVA with Tukey post-hoc). (I) Secreted lactate (ng µl⁻¹) in WT, BST2OE and BST2KO adipocytes at D6. ***p < 0.001 (one-way ANOVA with Tukey post-hoc). (J) Mass-isotopologue distributions (m+0 through m+6) for seven glycolytic and glycerol-3-phosphate metabolites after a 24-h pulse of uniformly ¹³C-labelled D-glucose in WT and BST2KO adipocytes. Bars are mean ± s.e.m. (n = 3 biological replicates).

To test whether BST2 controls adipocyte metabolism, we performed RNA-seq on WT, BST2KO (*CRISPR* KO) and BST2OE (stable overexpression) 10T1/2 adipocytes (adipocytes) ± norepinephrine (NE) stimulation (N=3/condition). Differential expression across total 12 conditions revealed a reciprocal glycolytic signature: *Hk2, Pfkl, Pfkp, Aldoa, Gapdh, Pgk1, Pgam1, Eno1, Pkm* and *Tpi1* were coordinately induced in BST2OE and downregulated in BST2KO, both at baseline and under NE (Fig. 1C). A gene-level t-SNE of 12,038 expressed genes colored by mean log₂FC(OE − KO) revealed a discrete island of glycolytic enzymes that segregated along the BST2-direction axis, and the marginal density of glucose-metabolism genes was strongly right-shifted relative to all other genes (Kolmogorov–Smirnov D = 0.192, p = 5.79 × 10⁻⁷⁰; Mann–Whitney p = 2.26 × 10⁻⁷⁷; Fig. 1D,1E). Gene Ontology enrichment recapitulated this asymmetry: glucose- and glycogen-catabolic terms were positively enriched in BST2OE (NES = 2.10–4.40, FDR < 0.05) and negatively enriched in BST2KO across both control and NE conditions (Fig. S1H).

Targeted LC–MS/MS metabolomics on D6 differentiated adipocytes demonstrated that BST2KO adipocytes markedly accumulated intracellular glucose but showed decreased upper-glycolytic intermediates (G-6-P, F-6-P, F-1,6-BP, 2-PGA, 3-PGA, PEP) and pyruvate and lactate alongside increased amino acid pools Fig. 1F). Functionally, Seahorse extracellular-acidification-rate (ECAR) recapitulated this glycolytic defect (Fig. 1G, S1I). WT adipocytes responded to sequential injections of glucose, oligomycin and NE with the canonical glycolytic-stress profile, while BST2KO adipocytes showed minimal ECAR response in any condition (Fig. 1G; replicated in an independent cohort in Fig. S1I). Independent assays showed that G-6-P and lactate were elevated in BST2OE and depleted in BST2KO adipocytes (p < 0.001; Fig. 1H,I). The pool-size signature (Fig. 1F), indicative of a stalled pathway, was orthogonally confirmed by stable-isotope tracing with uniformly ¹³C-labelled glucose. Steady-state m+6 D-glucose retention rose ∼3-fold in BST2KO, while m+6 and m+3 labelling of downstream glycolytic species (F-6-P, F-1,6-BP, lactic acid, pyruvic acid, G-6-P) and the lipogenic branch metabolite glycerol-3-phosphate markedly decreased (Fig. 1J). Integration of pool size, fractional enrichment and total labelled exchange rate in a single heatmap (Fig. S1J–L) revealed a coherent system where in BST2KO, upper-glycolytic pool sizes and exchange rates decrease (e.g., pyruvate exchange rate −3.7%, ; F-1,6-BP exchange rate −2.2%; α-KG exchange rate −4.6%,), with concomitant increase in cellular glucose levels and in some of the components of pentose-phosphate, nucleotide, and amino-acid fluxes (Fig. S1J–L).

Together, the transcriptome, steady-state metabolome, ¹³C flux and bioenergetic data converge on a model in which BST2 is cell-autonomously required to sustain glycolytic throughput, and its loss in adipocytes phenocopies the obesity-associated suppression of BST2 observed in human and mouse adipose atlases.

### BST2 controls adipocyte lipid composition and MAVS–TBK1–IRF3 innate-immune signaling

Because BST2 is an ISG and its loss reshaped central carbon metabolism, we asked whether BST2 also gates lipid composition and innate-immune signaling. Shotgun lipidomics across WT, BST2KO and BST2OE adipocytes revealed broad lipid and triacylglycerol (TG) remodeling (Fig. 2A,B; Fig. S2A,B), with BST2OE elevating total TG ∼1.8-fold and BST2KO reducing it ∼2.3-fold relative to WT (Fig. S2D). Class-resolved analysis (Fig. 2C; Fig. S2C) revealed that cholesterol esters (CE), diacylglycerols (DAG) and free fatty-acid (FFA) species were selectively upregulated in BST2OE and downregulated BST2KO adipocytes, and inflammatory lipids such as ceramides and lactosylceramide (LacCer) were lower in BST2OE adipocytes. The top differentially high and low abundant species in each genotype, including specific HexCER, LPC, LPE, LacCER, PA, PC, PE, PE P-, PG, PI, PS and SM lipids, are shown in Fig. S2C,E.

**Figure 2.**
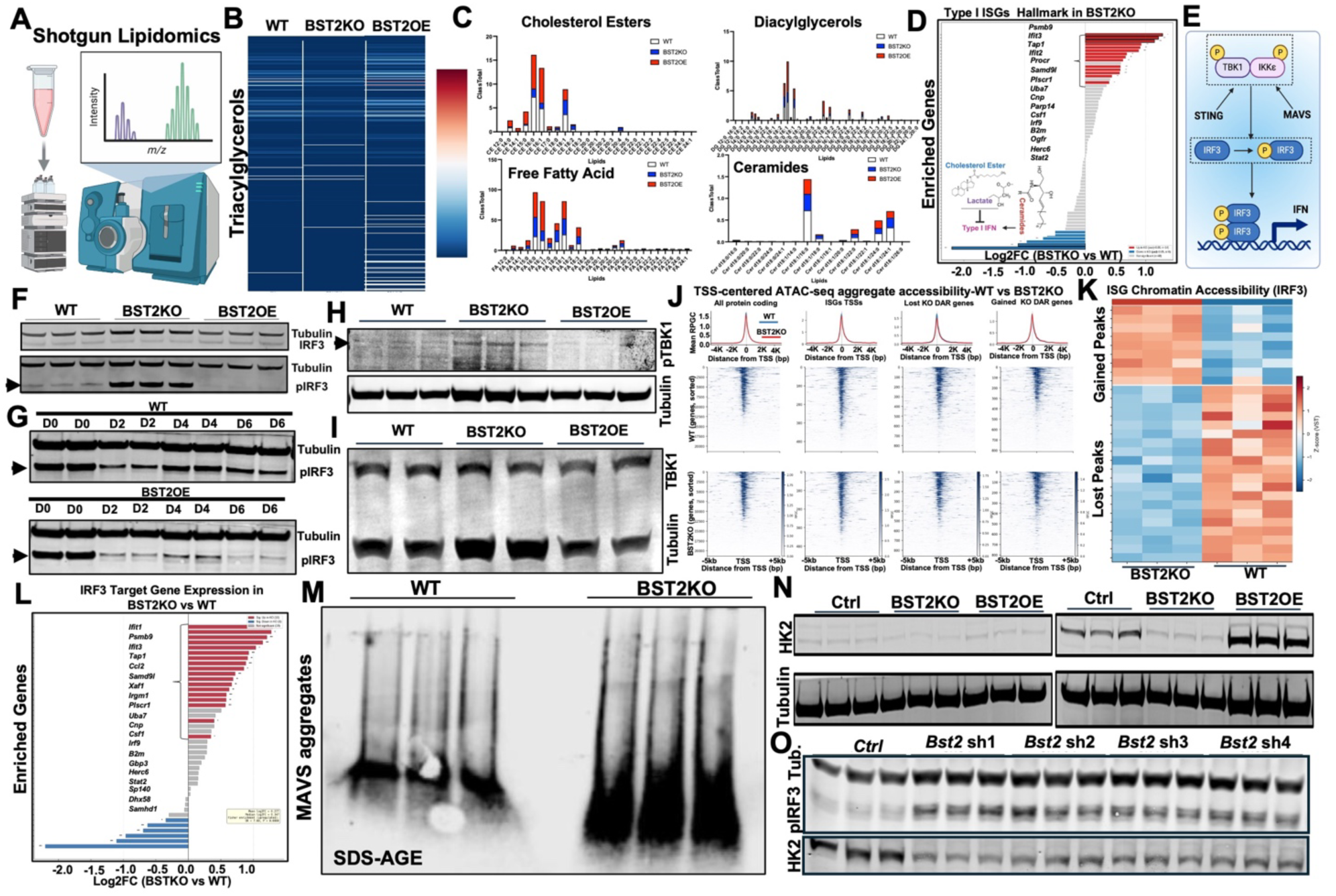
BST2 controls adipocyte lipid composition, MAVS–TBK1–IRF3 innate-immune signaling and chromatin accessibility. (A) Schematic of the shotgun-lipidomics workflow used to profile WT, BST2OE and BST2KO adipocytes. (B) Triacylglycerol species heatmap (z-scored intensities) across WT, BST2KO and BST2OE. (C) Class-resolved relative abundance for cholesterol esters, diacylglycerols, free fatty acids and ceramides across genotypes. Bars are stacked mean abundances. (D) Hallmark Type I ISG enrichment in BST2KO versus WT: ranked log₂FC bar chart of the top ISG leading-edge genes. Inset highlights cholesterol-ester and lactate, and ceramide species as candidate STING/MAVS-IRF3-IFN modulators. (E) Schematic of the canonical IFN ssignaling cascade. (F) Western blot of total IRF3 (IRF3), pIRF3 and Tubulin in WT, BST2KO and BST2OE adipocytes. (G) pIRF3 / Tubulin time-course (D0, D2, D4, D6) in WT and BST2OE adipocytes. (H) pTBK1 / Tubulin in WT, BST2KO and BST2OE adipocytes. (I) TBK1 / Tubulin in the same samples. (J) TSS-centered ATAC-seq aggregate accessibility (mean RPGC ± window, ±4 kb) in WT (top, blue) versus BST2KO (bottom, red), across four sets: all protein-coding genes, ISG TSSs, KO-lost differentially accessible regions (DARs), and KO-gained DARs. Heatmaps show per-gene accessibility sorted by WT signal. (K) ISG (IRF3-target) chromatin-accessibility heatmap with gained and lost peaks in BST2KO versus WT (z-score, log₂FC). (L) IRF3 target-gene expression bar chart (log₂FC, BST2KO vs WT) for the leading IRF3-target geneset; red = up, blue = down. (M) SDD-AGE native gel for MAVS aggregates in WT versus BST2KO adipocytes (n = 3 per genotype). (N and O) Western blot of HK2, pIRF3 and Tubulin in Ctrl, BST2KO, BST2OE (N) and four independent Bst2 shRNA knock-down lines (sh1–sh4) (O). Bars are mean ± s.e.m. (n = 3 biological replicates).

CE, ceramides, and lactate are known to be endogenous modulators of type I interferon signaling^46–49^. Hallmark gene-set enrichment showed a Type I ISG signature elevated in BST2KO adipocytes (Fig. 2D), and we investigated this signature on the canonical STING/MAVS: TBK1/IKKε:IRF3:IFN cascade (Fig. 2E). IRF3 is activated by phosphorylation at Ser 386^50,51^ (pIRF3) and phospho-IRF3 (pIRF3) was markedly elevated in BST2KO adipocytes, paralleled by elevated phospho-TBK1, while BST2OE markedly suppressed pIRF3 (Fig. 2F–I, Fig. S2F). As shown in Fig. 2F,G, levels of pIRF3 showed a drastic decrease during differentiation into adipocytes, and knockdown of BST2 rescues while BST2OE accelerates this downregulation. The same pattern was reproduced under poly(I:C) (polyinosinic: polycytidylic acid, a synthetic analog of double-stranded RNA (dsRNA)) challenge in Fig. S2G, suggesting that BST2 restricts Type I IFN signals and IRF3 phosphorylation/activation in adipocytes to potentially maintain immune and glycolytic balance^17,19^. In addition, ATAC-seq in WT versus BST2KO adipocytes mapped these gene-expression changes onto chromatin (Fig. 2J,K). TSS-centered aggregate accessibility was redistributed at protein-coding genes, ISG TSSs, and at sets of regions that gained or lost accessibility specifically in BST2KO. A focused heatmap of IRF3-target ISGs revealed coordinated gain-of-accessibility peaks paired with loss-of-accessibility peaks in BST2KO versus WT, consistent with active IRF3-driven remodeling (Fig. 2K). At the transcript level (Fig. 2L), classical IRF3 targets including *Ifit1, Psmb9, Ifit3, Tap1, Ccl2* and *Samd9l* were induced in BST2KO, whereas a subset of IRF3 targets co-regulated with metabolism were repressed. As shown earlier, both BST2KO and poly(I:C) treatment in adipocytes rescued the expression of pIRF3. Since dsRNA is known to activate MAVS-IRF3 pathway, our native SDD-AGE experiment confirmed that BST2KO adipocytes accumulated higher degree of MAVS aggregates, the structural correlate of active MAVS signaling^52^, compared to controls (Fig. 2M; Fig. S2L,M). Four independent *Bst2* shRNAs reproduced elevated pIRF3 and suppressed rate-limiting glycolytic enzyme HK2 in WT adipocytes, ruling out clonal artefacts of constitutive BSTKO cells (Fig. 2N,O). Joint RNA-seq/ATAC-seq concordance analysis (Fig. S2H–K) demonstrated that genes in the Glycolysis, FAO, TCA-cycle and Thermogenesis hallmark sets exhibited concordant RNA–chromatin behavior, anchoring metabolic gene-expression changes also in primary chromatin remodeling. Pathway enrichment of regions losing accessibility in BST2KO converged on fat-cell differentiation, cadherin/IgSF CAM signaling and morphogenesis (Fig. S2J), suggesting that BST2 loss simultaneously dampens adipose metabolic chromatin and enhances MAVS–IRF3 innate-immune program.

### Adipocyte-specific BST2 overexpression improves systemic glucose handling, reduces hepatic steatosis and increases energy expenditure in vivo

To test the in vivo consequences of altered adipocyte BST2 expression, we generated adipocyte-specific gain- and loss-of-function mouse models. The human adipocyte-specific BST2 overexpression (AdBST2OE) allele (CAG-loxP-Neo-Stop-loxP-hBST2-IRES-GFP-FRT-polyA) was driven by *Adipoq-Cre* to overexpress human BST2 selectively in mature adipocytes (AdhBST2OE), and the conditional Bst2ᶠ/ᶠ allele was crossed to Adipoq-Cre to ablate BST2 in mature adipocytes (AdBST2KO) (Fig. S3A). Real time qPCR confirmed transgene-driven hBST2 expression, BST2 knockdown, and levels of interferon (IFNA) and HK2 showed expected changes as seen in our in vitro system (Fig. S3A).

On a HFD, AdhBST2OE mice showed markedly improved glucose tolerance (AUC, p < 0.01) and insulin tolerance (AUC, p < 0.05) versus AdhBST2ᶠ/ᶠ littermates (Fig. 3A,B). Conversely, AdBST2KO mice exhibited markedly decreased glucose tolerance (total AUC, p < 0.05; Fig. 3C) and visibly elevated hepatic steatosis on H&E (Fig. 3D). A second AdhBST2OE cohort challenged with high sucrose diet (Western diet-WD) reproduced the AdhBST2OE driven improvement in both male and female mice glucose tolerance (AUC, p < 0.01; Fig. 3E,F), with histological evidence of reduced lipid accumulation (Fig. 3G,H) and a quantitatively lower hepatic steatosis index (∼10% vs ∼33%, p < 0.01; Fig. 3H). The AdhBST2OE phenotype extended to whole-body bioenergetics: 72-hour indirect-calorimetry data showed AdhBST2OE mice had significantly elevated energy expenditure (p < 0.01), lower fat oxidation (respiratory-exchange ratio) (p < 0.01) and increased oxygen consumption (p* < 0.01) across both light and dark phases (Fig. 3I–K).

**Figure 3.**
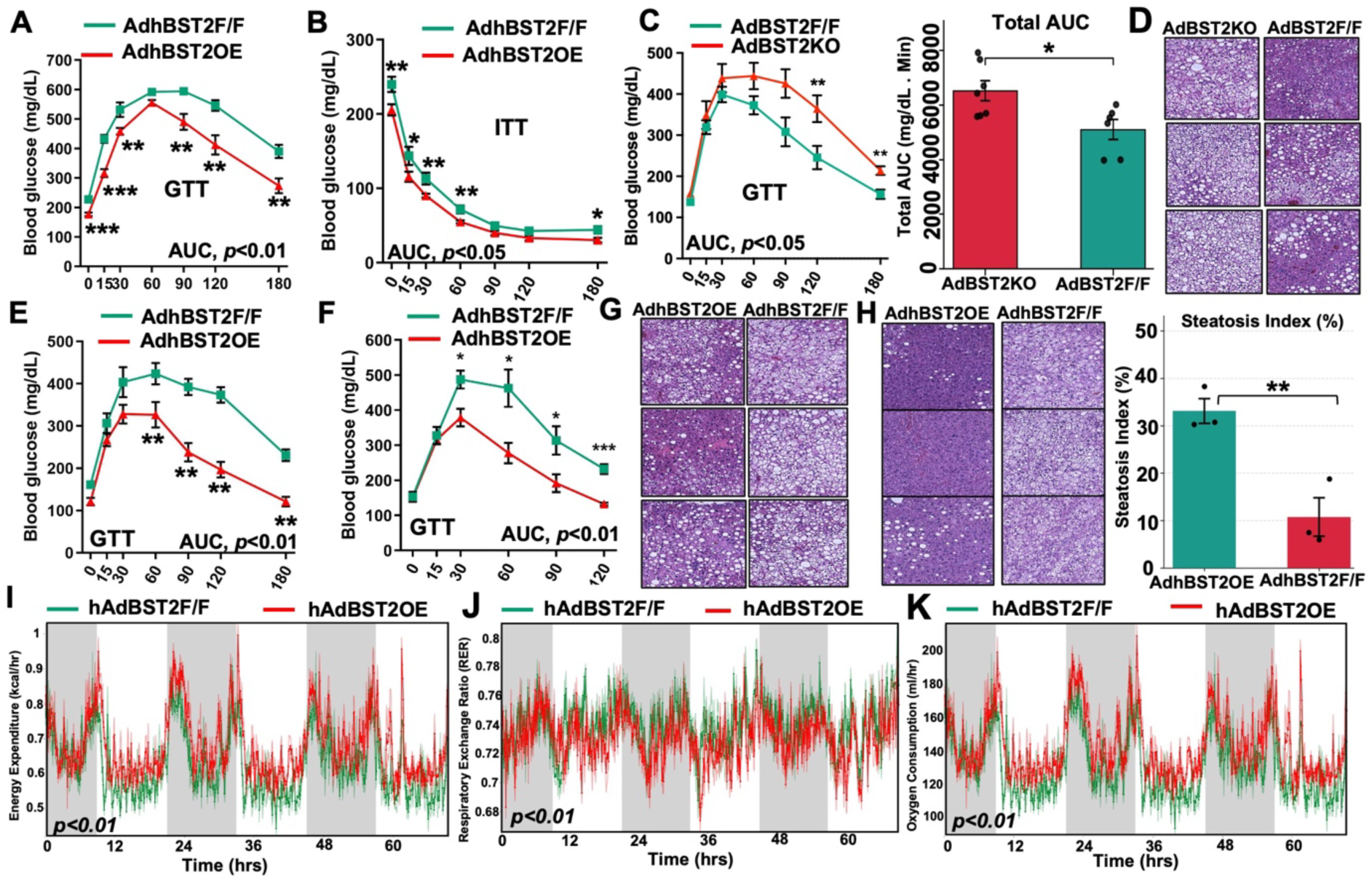
Adipocyte-specific BST2 overexpression improves systemic glucose handling and energy expenditure. (A) Glucose-tolerance test (GTT, 1 g kg⁻¹ i.p.) in AdhBST2ᶠ/ᶠ (green) versus AdhBST2OE (red) 12 weeks HFD-fed male mice. Blood glucose (mg dL⁻¹) vs time (min). AUC p < 0.01; time-point comparisons and **p < 0.0, ***p<0.001 (two-way ANOVA with Tukey post-hoc). N=8,8. (B) Insulin-tolerance test (ITT, 0.75 U kg⁻¹ i.p.) in the same cohort. AUC p < 0.05. and *p < 0.05, **p<0.01 (two-way ANOVA with Tukey post-hoc) N=8,8. (C) GTT in AdBST2KO versus AdBST2ᶠ/ᶠ 12 weeks HFD-fed male mice with total-AUC bar chart. AUC *p < 0.05. **p<0.01 (two-way ANOVA with Tukey post-hoc) N=7,8. (D) H&E histology of liver sections from AdBST2KO and AdBST2ᶠ/ᶠ mice (6 representative panels). (E and F) Independent GTT cohort from 14 weeks high sucrose diet fed male (E) and female (F) mice: AdhBST2OE versus AdhBST2ᶠ/ᶠ. AUC p < 0.01. **p<0.01 (two-way ANOVA with Tukey post-hoc) N=7,8. (G and H) H&E histology of HFD-FED (G) or high sucrose diet-fed (H) AdhBST2OE and AdhBST2ᶠ/ᶠ (A, B) liver sections. H&E histology with steatosis-index quantification (% area lipid). AdhBST2OE ∼10% vs AdhBST2ᶠ/ᶠ ∼33%, **p < 0.01. (I-K) Indirect calorimetry data showing energy expenditure (kcal h⁻¹) over 72 h with light/dark shading; respiratory-exchange ratio (RER) over 72 h; and oxygen consumption (VO₂, ml h⁻¹) over 72 h; p < 0.01 (ANCOVA) N=8,8. Bars are mean ± s.e.m.

Further metabolic characterization (Fig. S3B–I) demonstrated that AdhBST2OE mice gained largely comparable weight to controls across HFD and WD challenges in both sexes (Fig. S3B), but had reduced gWAT, iWAT and BAT depot masses (Fig. S3C), and reproduced the GTT/ITT improvement in females (Fig. S3D,E, AUC p < 0.01 and p < 0.05). Gross tissue images and histology (Fig. S3C, S3F–H) showed smaller adipocytes and reduced ectopic lipid in AdhBST2OE in BAT and liver. Protein HK2 levels were markedly increased in AdhBST2OE compared to controls (Fig. S3C). Female indirect calorimetry data (Fig. S3I) recapitulated the elevated VO₂ (p = 0.0041), energy expenditure (p = 0.0047) and RER shift (p = 0.00128) in AdhBST2OE. These data establish that white and brown adipocyte-intrinsic BST2 program, identified in vitro, improved systemic glucose homeostasis, attenuated hepatic steatosis and increased thermogenic energy expenditure in vivo.

### BST2 directly interacts with innate immune and glycolytic machinery to form a dual immunometabolic checkpoint in adipocytes

BST2KO and BST2OE failed to control IRF3 activation in preadipocytes compared to adipocytes (Fig. 4A), implicating a requirement of immunometabolic machinery that may be only present in adipocytes. BST2 was expressed in APCs and surface expression on PDGFRα⁺⁺ APCs was IFNAR1-dependent (Fig. S4A). Beige differentiated PDGFRα⁺⁺ BST2⁺⁺ cells were enriched for thermogenic markers *Ucp1, Ppargc1a* and *Cox8b* versus BST2⁻ APCs (Fig. S4B), however, preadipocyte-specific *Bst2* deletion (preAdBST2KO; *Pdgfra*-Cre × Bst2ᶠ/ᶠ) did not impair glucose tolerance compared to controls (Fig. S4C), suggesting the checkpoint operates from the adipocyte stage onward. To define the molecular machinery through which BST2 couples glycolysis to innate-immune signaling in adipocytes, we performed Flag-BST2 immunoprecipitation followed by quantitative LC–MS/MS in D6 10T1/2 differentiated adipocytes stable expressing Flag-BST2 (N = 3,3). Of proteins significantly enriched by Flag-BST2 over IgG, three were prioritized because they bridge the metabolic and immune aspects of the phenotype: the glycolytic feed-forward activator PFKFB3 (6-phosphofructo-2-kinase/fructose-2,6-bisphosphatase 3), the E3-ligase RNF128^53^ and the Nedd4-family interactor NDFIP1^54^ (Fig. 4B). Pairwise and trimeric AlphaFold/Boltz2^55,56^ modelling predicted BST2’s coiled-coil ectodomain as the interaction surface for RNF138 and cytoplasmic domain for PFKFB3 and NDFIP1, with confidence scores 0.56–0.70, pTM 0.40–0.59 and ipTM 0.21–0.47 across BST2–PFKFB3, BST2–RNF128 and BST2–NDFIP1 binary complexes and trimer combinations (Fig. 4C; Fig. S4H). Furthermore, all three BST2 interactors were present in adipocytes and were upregulated during differentiated adipocytes, and BST2 did not control their expression in adipocytes (Fig. S4D-S4F). Functional dissection of each node was performed in CRISPR-Cas9 clonal lines. NDFIP1 knockout (Fig. 4D, Cl. A–D) and RNF128 knockout (Fig. 4E, Cl. A–D) both stabilized adipocytes pIRF3, identifying RNF128/NDFIP1 as a BST2-dependent restraint on IRF3 activation.

**Figure 4.**
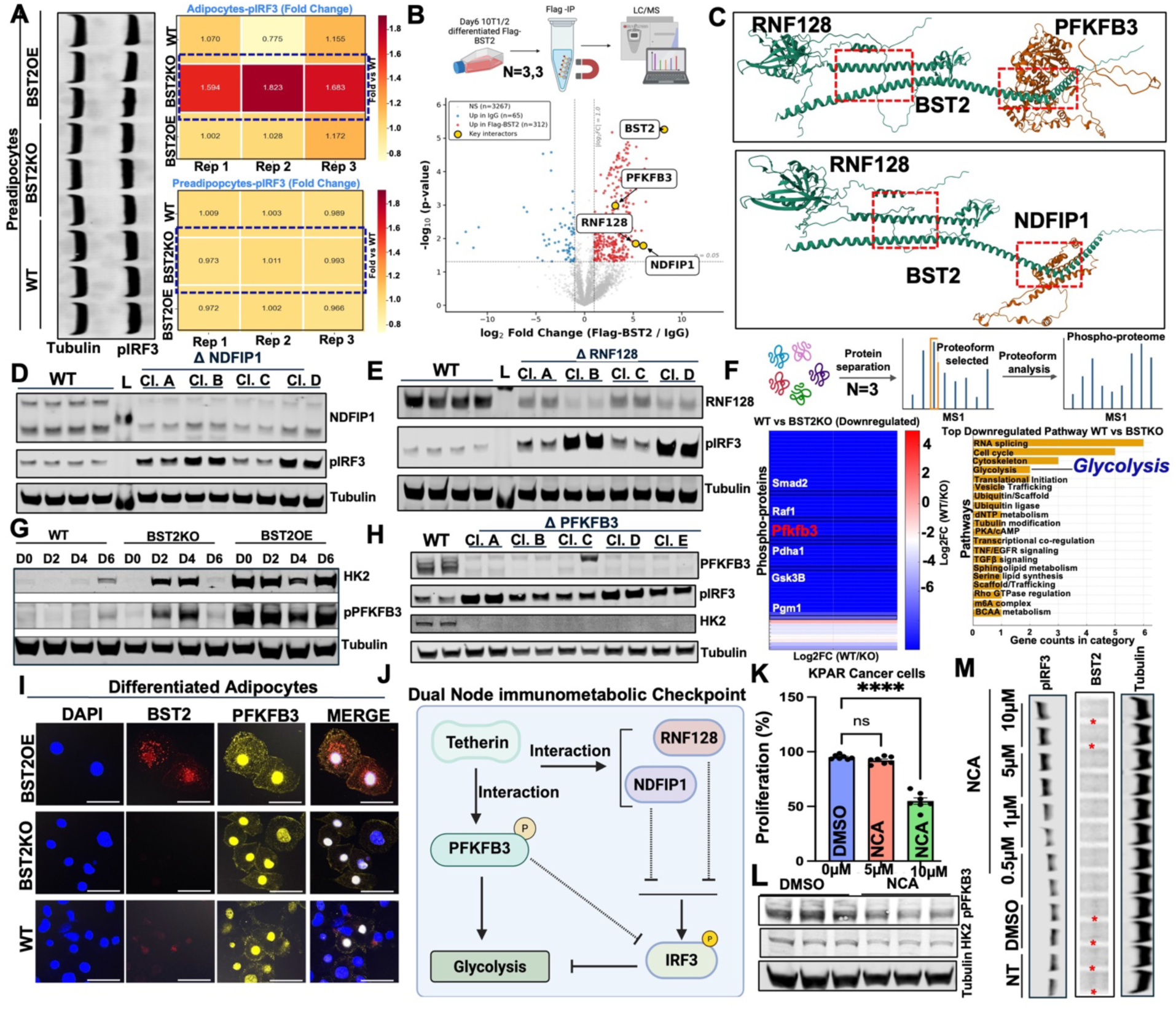
BST2 engages PFKFB3, RNF128 and NDFIP1 to form a dual pathway immunometabolic checkpoint. (A) Western blot of Tubulin and pIRF3 in preadipocytes and differentiated adipocytes (WT, BST2KO, BST2OE) with quantified fold-change heatmaps (3 replicates per condition). Highlighted dotted rectangles show pIRF3 is elevated in BST2KO adipocytes but not in preadipocytes. (B) Volcano plot of Flag-BST2 IP-MS in day-6 10T1/2 differentiated adipocytes (N = 3 vs 3). 312 proteins are significantly enriched in Flag-BST2 over IgG; key interactors BST2, PFKFB3, RNF128 and NDFIP1 are highlighted. (C) AlphaFold-predicted complexes of BST2 with RNF128, PFKFB3 and NDFIP1; predicted interaction interfaces are boxed in red dashed outlines. (D) Western blot of NDFIP1, pIRF3 and Tubulin in WT and four *CRISPR* NDFIP1-KO clones (Cl.A–D). (E) Western blot of RNF128, pIRF3 and Tubulin in WT and four *CRISPR* RNF128-KO clones (Cl.A–D). (F) Phospho-proteomics workflow (N = 3 combined) and heatmap of down-regulated phospho-proteins in BST2KO vs WT (Smad2, Raf1, Pfkfb3, Pdha1, Gsk3B, Pgm1); pathway-enrichment bar chart with Glycolysis among the top down-regulated pathways. (G) Time-course Western blot (D0, D2, D4, D6) for HK2, pPFKFB3 and Tubulin in WT, BST2KO and BST2OE adipocytes. (H) Western blot of PFKFB3, pIRF3, HK2 and Tubulin in WT and *CRISPR* PFKFB3-KO clones (Cl.A–E). (I) IF of differentiated BST2OE, BST2KO and WT adipocytes stained for DAPI (blue), BST2 (red), PFKFB3 (yellow) and MERGE. Scale bar, 100 µm. (J) Schematic of the Dual-Node Immunometabolic Checkpoint: Tetherin/BST2 engages PFKFB3 to support glycolysis and engages RNF128/NDFIP1 to restrain pIRF3, with reciprocal cross-regulation between the two arms. (K) KPAR proliferation (%) under DMSO, 5 µM NCA and 10 µM NCA. ****p < 0.0001 (one-way ANOVA). (L) Western blot of pPFKFB3, HK2 and Tubulin in KPAR cells treated with DMSO or 10 µM NCA. (M) Western blot of pIRF3, BST2 and Tubulin in KPAR cells across a NCA dose series (NT, DMSO, 0.5, 1, 5, 10 µM). Bars are mean ± s.e.m. (n = 3 biological replicates).

Unbiased phospho-proteomics (Fig. 4F) in WT versus BST2KO adipocytes (N = 3 combined/condition) demonstrated markedly attenuated phosphorylation of glycolytic regulators (Pfkfb3, Pgm1, Pdha1) and signaling kinases (Smad2, Raf1, Gsk3B) in BST2KO, with Glycolysis identified as a top down-regulated pathway alongside RNA splicing, cell cycle, cytoskeleton and translation. This was further confirmed by western blot where time-course analysis across D0–D6 of differentiation showed that PFKFB3 phosphorylation (pPFKFB3 at Ser 461)^57–59^ was markedly elevated with differentiation in BST2OE D6 adipocytes, while HK2 protein showed the same pattern (Fig. 4G). Furthermore, PFKFB3 knockout decreased adipocyte HK2 protein (Fig. 4H), placing PFKFB3 epistatically upstream of HK2 protein abundance, and markedly increased pIRF3 levels. However, 2-deoxyglucose (2DG) blockade of glycolysis still suppressed pIRF3 in BST2OE adipocytes, suggesting that BST2 restraint of innate immune activation did not rely on activating glycolytic pathway (Fig. S4G). Immunofluorescence in differentiated adipocytes confirmed BST2–PFKFB3 co-staining and increased cytoplasmic PFKFB3 levels in BST2OE cells with loss of signal in BST2KO (Fig. 4I; not in preadipocyte in Fig. S4I), suggesting an interaction-dependent spatial cytoplasmic retention/organization of PFKFB3 for potential activation (phosphorylation, pPFKFB3) by kinases to control glycolysis^58–61^ (Fig. S4J). Overall, we integrated these data into a Dual-Node Immunometabolic Checkpoint model: BST2 (Tetherin) at the ER simultaneously engages PFKFB3 to control glycolysis and engages RNF128/NDFIP1 to restrain pIRF3, with glycolysis and IRF3^17^ reciprocally cross-regulated (Fig. 4J), establishing a feed-forward loop between BST2-controlled type I interferon signaling and glycolysis.

We next asked whether BST2’s immunometabolic role is conserved in other metabolically active cells. Across the TCGA pan-cancer cohort (n = 9,295)^62^, BST2 expression correlated positively with the Hallmark Glycolysis signature (Pearson r = 0.243; Spearman ρ = 0.224; p < 1e-100; Fig. S4K), with significant per-cancer concordance in 25 of ∼33 tumor types (FDR < 0.05; Fig. S4L). In TCGA-PAAD (pancreatic adenocarcinoma, n = 178), BST2 correlated with HK2 (r = 0.471, p = 3.0 × 10⁻¹¹) and PFKFB3 (r = 0.342, p* = 2.6 × 10⁻⁶; Fig. S4M), and the BST2–glycolysis coupling held across lung-adenocarcinoma KRAS subgroups ^62^ (Fig. S4N) and across independent T1D/T2D pancreatic cohorts^63–66^ (Fig. S4O). We treated KPAR lung cancer cells with the selective BST2 inhibitor, Neocarzilin (NCA)^67^. NCA dose-dependently suppressed KPAR proliferation (** p < 0.0001 at 10 µM; Fig. 4K), reduced HK2 and pPFKFB3, and concurrently increased pIRF3 (Fig. 4L,M; reproduced for HK2 and pFKFB3 in INS1 insulinoma cells across three independent biological sets under high-glucose and cytokine-cocktail conditions in Fig. S4P). Thus, the BST2–PFKFB3–IRF3 checkpoint is a conserved, pharmacologically relevant axis that operates in both immunometabolic and oncogenic glycolytic settings.

## Discussion

Our study identifies BST2 as an adipocyte immunometabolic checkpoint that links resolution of innate immune signaling to preservation of glycolytic capacity. Although BST2 is classically known as an interferon-inducible antiviral restriction factor, our findings support a broader physiological role in metabolic cells. In adipocytes, BST2 does not simply mark an inflammatory state; rather, it actively organizes a cell-intrinsic program that suppresses IRF3-driven interferon signaling while sustaining core glycolytic machinery, including PFKFB3 phosphorylation and HK2 abundance. This dual function places BST2 at a mechanistic intersection between immune-state control and metabolic homeostasis.

A central conceptual advance of this work is the reframing of inflammatory signaling and metabolic dysfunction as directly coupled phenomenon in adipocytes. Instead of treating inflammation and metabolism as parallel outputs, our data support the idea that unresolved antiviral-like signaling in adipocytes can be a proximal driver of metabolic failure. Loss of BST2 increased pIRF3, induced interferon-responsive genes, altered lipid composition, and reduced glycolytic intermediates and flux. Our data further suggest that BST2 exerts this checkpoint function through coordinated control of both signaling and metabolism. On the immune side, BST2 interacted with RNF128 and NDFIP1, and loss of either factor increased IRF3 activation, consistent with a model in which BST2 assembles or stabilizes a signal-terminating complex that restrains MAVS–TBK1–IRF3 activity. On the metabolic side, BST2 associated with PFKFB3, promoted a phosphorylation active state, and was linked to a higher cytoplasmic retention of PFKFB3, a localization pattern consistent with enhanced glycolytic function. These observations support a model in which BST2 is not merely an upstream suppressor of inflammation or a parallel regulator of metabolism, but instead couples the two processes through discrete molecular interactions. In this framework, BST2 enables adipocytes to remain metabolically active while avoiding maladaptive antiviral stress program.

The adipocyte differentiation stage specificity of the phenotype is also notable. BST2-dependent differences in pIRF3 were not apparent in preadipocytes but emerged in mature adipocytes, indicating that BST2 function is context dependent and likely linked to the specialized organellar architecture and metabolic demands of the differentiated adipocytes. This is consistent with our imaging data placing BST2 in ER-associated compartments, where it would be well positioned to sense or regulate membrane, lipid-droplet, and signaling dynamics. Because differentiated adipocytes are uniquely programmed with buffering nutrient excess while maintaining endocrine and bioenergetic function, they may be especially dependent on checkpoint systems that prevent persistent innate activation from highly demanding metabolic programs for forming active fat cells.

The in vivo findings establish that BST2 cell-autonomous pathway has organismal metabolism relevance. Adipocyte BST2 gain-of-function improved glucose tolerance and insulin sensitivity, increased energy expenditure, and reduced hepatic steatosis under dietary stress, whereas adipocyte BST2 loss impaired glucose homeostasis. These reciprocal phenotypes indicate that BST2 is a major modulator of adipocyte state and systemic metabolic health. Our data also suggests that preserving adipocyte immunometabolic integrity can secondarily protect distal organs from ectopic lipid deposition and metabolic dysfunction. This positions adipocyte BST2 as a potential upstream node controlling whole-body energy partitioning during nutritional stress.

Our findings also have broader implications beyond adipose biology. In lung cancer cells, pharmacologic degradation of BST2 reduced proliferation while increasing pIRF3 and altering glycolytic signaling, suggesting that the BST2-centered coupling of immune restraint and metabolic support may extend to other metabolic cell types. Although the metabolic requirements and signaling context of cancer cells differ from those of adipocytes, both settings may benefit from a BST2-dependent state that suppresses innate stress signaling while preserving glycolytic output. This orthogonal evidence strengthens the general principle that BST2 can function as an immunometabolic organizer rather than solely as a viral tethering factor. It also raises the possibility that BST2 may be targeted differently across diseases: enhancing its activity in metabolic disease to preserve tissue homeostasis, while inhibiting it in cancer to destabilize tumor growth and metabolism. Finally, broad immunosuppression is neither mechanistically precise nor clinically ideal, and our finding opens a possibility of investigating specific cell intrinsic/extrinsic immune-metabolic checkpoints to better understand and treat metabolic diseases.

## Materials and Methods

### Cell culture and adipocyte differentiation

Mouse 10T1/2 cells (CCL-226, ATCC) were maintained in Dulbecco’s modified Eagle’s medium (DMEM; MT10013CM, Corning) supplemented with 10% fetal bovine serum (FBS; FB-11, Omega Scientific) and 1% penicillin–streptomycin (MT30002CI, Corning) at 37 °C in a humidified incubator with 5% CO₂. For routine maintenance, cells were washed with PBS (BP39920, Fisher Scientific) and detached using trypsin–EDTA (MT25053CI, Corning). Cells were passaged before reaching full confluence and confirmed to be free of mycoplasma contamination. For adipocyte differentiation, 10T1/2 cells were seeded at 50,000 cells ml⁻¹ in 6-well plates (07-200-83, Corning) with 2 ml of pre-adipogenic medium containing DMEM, 10% FBS, 1 nM T3 (T2877, Sigma-Aldrich) and 5 μg ml⁻¹ insulin (12-585-014, Gibco), and grown to confluence. Differentiation was initiated on day 0 by replacing the medium with adipogenic induction medium containing DMEM, 10% FBS, 1 nM T3, 5 μg ml⁻¹ insulin, 2 μg ml⁻¹ dexamethasone (D1756, Sigma-Aldrich), 0.5 mM 3-isobutyl-1-methylxanthine (IBMX; I7018, Sigma-Aldrich), 1 μM rosiglitazone (R2408, Sigma-Aldrich) and 125 μM indomethacin (I7378, Sigma-Aldrich), unless otherwise indicated. After 48 h, cells were switched to maintenance medium containing DMEM, 10% FBS, T3, insulin and rosiglitazone. Maintenance medium was replaced every 2 days. Cells were collected at the indicated differentiation stages for immunoblotting, RNA extraction or immunofluorescence microscopy. Where indicated, cells were stimulated with norepinephrine, CL-316,243 (C5976, Sigma-Aldrich) or DMSO vehicle control (J66650.AK, Thermo Fisher Scientific) at the concentrations and durations specified in the figure legends.

### KPAR cell culture and BST2 degrader treatment

KPAR lung cancer cells were provided by Dr. Daniel Puleston at the Icahn School of Medicine at Mount Sinai. Cells were cultured in Dulbecco’s modified Eagle’s medium (DMEM) supplemented with 10% fetal bovine serum and 1% penicillin–streptomycin. Cells were maintained at 37 °C in a humidified incubator with 5% CO₂ and passaged before reaching full confluence. For routine maintenance, cells were washed with PBS and detached using trypsin–EDTA. Cells were routinely monitored for morphology and growth rate and confirmed to be free of mycoplasma contamination. For BST2 degrader experiments, KPAR cells were seeded in 6-well plates for immunoblotting or in 96-well plates (07-200-90, Corning) for proliferation assays. After overnight attachment, cells were treated with DMSO vehicle or the BST2 degrader NCA (HY-125900, MedChemExpress) at the indicated concentrations. NCA was prepared as a stock solution in DMSO and diluted into complete culture medium immediately before treatment. The final DMSO concentration was kept consistent across all treatment groups. Cells were collected at the indicated time points for immunoblotting or proliferation assays.

### Proliferation assay

Cell proliferation was measured using crystal violet staining. Cells were seeded in 96-well plates at 2,000 cells per well and treated with DMSO vehicle or NCA at the indicated concentrations. At the experimental endpoint, culture medium was removed and cells were gently washed with PBS. Cells were fixed with 4% paraformaldehyde for 15 min at room temperature and stained with 0.1% crystal violet solution for 15 min. Plates were washed extensively with water to remove excess dye and air-dried completely. Bound crystal violet was solubilized with 10% acetic acid (695092, Sigma-Aldrich), and absorbance was measured at 590 nm using a plate reader. Values were normalized to the DMSO-treated control group.

### INS-1 cell culture

The rat insulinoma β-cell line INS-1 was cultured in RPMI 1640 medium (11875093, Gibco) supplemented with 10% fetal bovine serum, 1% penicillin–streptomycin, 2 mM L-glutamine (25030081, Gibco), 10 mM HEPES (15630080, Gibco), 1 mM sodium pyruvate (11360070, Gibco) and 50 μM β-mercaptoethanol (31350010, Gibco). Cells were maintained at 37 °C in a humidified incubator with 5% CO₂ and passaged before reaching full confluence. For routine maintenance, cells were washed with PBS and detached using trypsin–EDTA. Cells were routinely monitored for morphology and growth rate and confirmed to be free of mycoplasma contamination. For experiments, INS-1 cells were seeded at 600,000 cells per well in 6-well plates and allowed to attach overnight. Where indicated, cells were exposed to altered glucose conditions, cytokine cocktail treatment and/or BST2 degrader treatment for 72 h. For high-glucose experiments, cells were cultured under baseline glucose conditions of 11 mM and then switched to medium containing 20 mM glucose using D-glucose solution (A2494001, Gibco). For inflammatory stress experiments, cells were treated daily for 3 h with a cytokine cocktail consisting of 500 U ml⁻¹ interleukin-1β (IL-1β; 501-RL, R&D Systems), 1,000 U ml⁻¹ tumour necrosis factor-α (TNF-α; 510-RT, R&D Systems) and 100 U ml⁻¹ interferon-γ (IFN-γ; 585-IF, R&D Systems). For BST2 degrader experiments, cells were treated daily with DMSO vehicle or 10 μM NCA. Cells were harvested at the indicated time points for downstream immunoblotting analysis.

### Adenoviral BST2 knockdown in 10T1/2 cells

The adenoviral mouse Bst2 shRNA construct targeted the sequence 5′-GGGTTACCTTAGTCATCCTGA-3′. The full shRNA insert sequence was 5′-CACCGGGTTACCTTAGTCATCCTGACGAATCAGGATGACTAAGGTAACCC-3′. Adenoviruses were packaged and produced in 293A cells, and viral titres were determined by plaque assay. Mouse 10T1/2 cells were differentiated into adipocytes as described above. For adenoviral knockdown experiments, fully differentiated adipocytes were infected with adenovirus expressing mouse Bst2-targeting shRNA or Ad-mCherry as a control. The adenoviruses were provided by Dr. Peng Wang at the Icahn School of Medicine at Mount Sinai. Before infection, cells were gently washed once with PBS and incubated in serum-free DMEM without fetal bovine serum or antibiotics. Adenovirus was added at 2 × 10⁸ or 2 × 10⁹ PFU per well in 1 ml serum-free DMEM and incubated with cells for 2 h at 37 °C in a humidified incubator with 5% CO₂. After infection, the viral medium was removed and replaced with fresh adipocyte maintenance medium containing serum. Cells were maintained for 72 h after infection before collection for downstream analysis. BST2 knockdown efficiency was assessed by quantitative PCR and/or immunoblotting.

### Gene expression analysis

Total RNA was extracted from inguinal white adipose tissue using TRIzol reagent (15596026, Invitrogen) according to the manufacturer’s protocol. RNA quality and concentration were assessed by spectrophotometry. cDNA was synthesized from 500 ng of total RNA using the High-Capacity cDNA Reverse Transcription Kit (4374967, Applied Biosystems) according to the manufacturer’s instructions. Quantitative real-time PCR was performed using a QuantStudio 5 Real-Time PCR System (Applied Biosystems) in 384-well plates (4309849, Applied Biosystems) with Universal SYBR Green Fast qPCR Mix (RM21203, ABclonal). Gene expression was quantified using the standard curve method. All samples were analyzed in quadruplicate technical replicates, and expression levels were normalized to the housekeeping gene 36B4/Rplp0. Primer sequences are provided in Supplementary Table 1. The qPCR cycling conditions were as follows: initial denaturation at 95 °C for 10 min, followed by 40 cycles of denaturation at 95 °C for 15 s and annealing/extension at 60 °C for 30 s. Melting curve analysis was performed to confirm amplification specificity.

### Immunofluorescence microscopy

Cells were grown in black 96-well imaging plates (165305, Thermo Fisher Scientific), washed with PBS and fixed with 4% paraformaldehyde (AAJ19943K2, Thermo Fisher Scientific) for 15 min at room temperature. After fixation, cells were washed with PBS and permeabilized with 0.005% digitonin (D141, Sigma-Aldrich) in PBS for 10 min. Cells were then blocked for 1 h at room temperature in blocking buffer containing 10% normal goat serum (NGS; ab7481, Abcam), 1% bovine serum albumin (BSA; A4737, Sigma-Aldrich) and 0.001% digitonin in PBS. Primary antibodies were diluted in blocking buffer and incubated overnight at 4 °C. White adipose tissues were collected from mice under the indicated experimental conditions and fixed overnight in 10% neutral buffered formalin (50-134-4524, G-Biosciences). After fixation, tissues were washed with PBS and stored in 70% ethanol (BP28184, Fisher Scientific) at 4 °C until processing. Paraffin embedding and sectioning were performed by the Mount Sinai Histology Core. Tissue sections were deparaffinized by sequential 10-min incubations in xylene twice (X3P, Fisher Scientific), 100% ethanol twice, 95% ethanol twice, 85% ethanol twice and 75% ethanol twice. Slides were then washed three times with water before antigen retrieval.

For antigen retrieval, slides were immersed in pre-heated Tris–EDTA buffer containing 10 mM Tris base and 1 mM EDTA. Slides were heated in a microwave at 98 °C for 20 min and then cooled in cold tap water for 10 min. Tissue sections were permeabilized with 0.05% Tween-20 (655205, Millipore) in PBS for 10 min at room temperature. Sections were blocked for 1 h at room temperature in blocking buffer containing 10% NGS, 1% BSA and 0.05% Tween-20 in PBS. Primary antibodies were diluted in blocking buffer and incubated overnight at 4 °C. After primary antibody incubation, cell and tissue samples were washed and incubated with species-appropriate fluorophore-conjugated secondary antibodies for 1 h at room temperature protected from light. Nuclei were counterstained with DAPI (EN62248, Invitrogen) for 5 min at room temperature. Cells were washed and imaged in PBS, whereas tissue sections were mounted using antifade mounting medium (P10144, Invitrogen). Images were acquired using a Leica SP8 AOBS confocal microscope at the Mount Sinai Microscopy Core. Laser power, detector gain and exposure settings were kept constant for comparisons within each experiment. Images were processed using Fiji/ImageJ.

### Western blot

Cells or tissues were lysed in ice-cold RIPA lysis buffer (NC9923398, Boston BioProducts) supplemented with protease and phosphatase inhibitors (PIA32961PM, Thermo Fisher Scientific). Lysates were incubated on ice for 5 min and clarified by centrifugation at 13,000 × g for 15 min at 4 °C. Protein concentrations were determined using a BCA assay (PI23227, Thermo Fisher Scientific). Equal amounts of protein were mixed with LDS sample buffer (B0007, Invitrogen) containing 50 mM dithiothreitol (DTT; FERR0861, Thermo Fisher Scientific), denatured at 100 °C for 15 min, separated by SDS–PAGE and transferred to PVDF membranes (IPFL85R, Sigma-Aldrich). Membranes were blocked with LI-COR blocking buffer (927-60010, LI-COR) and incubated with primary antibodies overnight at 4 °C. After washing, membranes were incubated with fluorescent secondary antibodies for 1 h at room temperature. Signals were detected using a LI-COR Odyssey imaging system, and band intensities were quantified using Empiria Studio. Protein abundance was normalized to tubulin or to the corresponding total protein, as indicated in the figure legends. All antibodies used are listed in Supplementary Table 2.

### Indirect calorimetry

Whole-body energy metabolism was assessed using an indirect calorimetry system (Promethion Core, Sable Systems). Mice were individually housed in metabolic cages under controlled temperature (22 °C), humidity (50%) and a 12-h light–dark cycle, with lights on from 7:00 a.m. to 7:00 p.m. Mice had ad libitum access to food and water unless otherwise stated. Mice were acclimated to the metabolic cages for 72 h before data collection. Oxygen consumption, carbon dioxide production, respiratory exchange ratio, energy expenditure, locomotor activity and food intake were measured continuously at 5-min intervals. Data from the acclimation period were excluded from analysis. Light- and dark-cycle values were analyzed separately where appropriate. Energy expenditure was analyzed using body weight.

### TCA Metabolomics

#### Metabolite extraction

Metabolites were extracted from 10T1/2 cells differentiated for 6 days in 6-well plates. Cells were washed, and extraction was initiated by adding 1 mL of ice-cold 80:20 (v/v) methanol:water. Samples were lysed using ceramic bead tubes with 10 cycles of 15 s shaking at 600 m/s on an Omni Bead Ruptor homogenizer. Lysates were centrifuged at 20,000 × g for 20 min at 4°C, and the resulting supernatants were transferred to fresh tubes. Extracts were dried under vacuum using a Genevac EZ-2 Plus evaporator for 2 h. Dried samples were reconstituted in 50 μL of 50% (v/v) acetonitrile, vortexed, and incubated on ice for 20 min, followed by clarification by centrifugation at 20,000 × g for 20 min at 4°C prior to LC-MS analysis.

#### Metabolite Detection by LC-MS/MS

Metabolite detection was carried out using HILIC chromatography with an UHPLC system (Vanquish Horizon, Thermo Fisher Scientific) coupled to an Orbitrap IQ-X Tribrid mass spectrometer (Thermo Fisher Scientific) operated in heated electrospray ionization (HESI) mode. Chromatographic separation was performed on a Waters Atlantis Premier BEH Z-HILIC column (2.1 × 150 mm, 1.7 μm). A 2 μL sample injection volume was used, with the column oven maintained at 30°C and a flow rate of 0.2 mL/min. Mobile phase A consisted of water with 10 mM ammonium carbonate, and mobile phase B consisted of 95% acetonitrile with 5 μM medronic acid (InfinityLab Deactivator Additive, Agilent). The gradient was as follows: 0–10 min, 80% to 50% B; 10–11 min, 50% to 80% B; 11–20 min, re-equilibration at 80% B. MS analyses were performed using a HESI source in negative polarity mode with the following source conditions: static spray voltage; sheath gas flow rate, 20 arbitrary units; aux gas flow rate, 4 arbitrary units; sweep gas flow rate, 3 arbitrary units; capillary temperature, 300°C; vaporizer temperature, 200°C; RF lens, 60%. Full-scan MS¹ spectra were acquired in the Orbitrap analyzer at a resolution of 240,000, with an AGC target of 4 × 10⁵, maximum injection time set to auto, and a scan range of 100–800 m/z in profile mode. Data analysis, including fragment ion extraction and peak area integration, was performed using Skyline-daily (v24.1.1.398). Stable isotope labeling analysis was performed using Compound Discoverer 3.4, where natural abundance correction was performed against samples cultured with unlabeled ^12^C glucose.

#### CRISPR–Cas9 genome editing in 10T1/2 cells

Single-cell knockout populations of 10T1/2 cells were generated by CRISPR–Cas9 ribonucleoprotein (RNP) electroporation. Cells were cultured in Dulbecco’s Modified Eagle Medium (DMEM) supplemented with 10% fetal bovine serum (FBS) and 1% penicillin–streptomycin (Thermo Fisher Scientific) at 37°C with 5% CO₂. Single guide RNAs (sgRNAs) targeting *Bst2*, *Ndfip1*, *Rnf128*, and *Pfkfb3* were designed in silico and synthesized by GenScript. For each target gene, multiple sgRNAs were pooled prior to RNP complex formation to maximize knockout efficiency (sgRNA sequences are listed in Supplementary Table 3). Recombinant Cas9 protein (TrueCut Cas9 Protein v2, Cat# A36498; Invitrogen) was complexed with the pooled sgRNAs at a 1.2:1 molar ratio in Resuspension Buffer R (Thermo Fisher Scientific) and incubated for 15 min at room temperature to form RNP complexes. Cells were detached using trypsin–EDTA (Invivogen), washed with DPBS (Gibco), and resuspended in Neon Resuspension Buffer R. Ten microliters of cell suspension was mixed with RNP complexes and electroporated using the Neon Transfection System (Thermo Fisher Scientific) at 1,300 V, 20 ms, and 2 pulses. Following electroporation, cells were immediately transferred into pre-warmed complete medium (DMEM with 10% FBS) and cultured under standard conditions, with medium replacement after 12–24 h. After sufficient expansion, genomic DNA was extracted using a standard lysis-based method; target loci were amplified by PCR and submitted for Sanger sequencing (Genewiz) to confirm indel formation and knockout status. For clonal isolation, cells were diluted and seeded at approximately one cell per well in 96-well plates (Corning). Individual clones were expanded prior to genomic DNA extraction and genotyping by PCR followed by Sanger sequencing to identify knockout clones.

### Immunoprecipitation and LC-MS/MS analysis

For LC-MS/MS analysis of BST2 interacting proteins in 10T1/2 differentiated cells, cell pellets were lysed using a HEPES-based lysis buffer (10 mM HEPES pH 7.4, 10 mM KCl, 0.05% NP-40) supplemented with protease and phosphatase inhibitors (Sigma-Aldrich, Cat# P8340; Fisher Scientific, Cat# A32957) for 30 min on ice. Homogenates were centrifuged at 16,000×g for 15 min at 4°C, and the supernatants were collected. Protein concentrations were determined using a BCA protein assay kit (Thermo Fisher Scientific, Cat# 23225). For antibody coupling, 3 mg of Dynabeads® M-270 Epoxy beads (Dynabeads® Antibody Coupling Kit; Fisher Scientific, Cat# 14311D) were coupled with 50 μg of either rat IgG isotype control (Cat# 31933; Thermo Fisher Scientific) or rat monoclonal anti-BST2 antibody (clone 120G8.04, Cat# DDX0390P-100; Novus Biologicals) per sample, and incubated for approximately 18 h at 37°C under continuous rotation. Antibody-coupled beads were washed sequentially and blocked with PBS containing 0.1% BSA prior to use. Lysates containing 2.5 mg of total protein were then incubated with the antibody-conjugated beads for 18–24 h at 4°C under continuous rotation to capture immune complexes. After sequential washes with bead washing buffers, immune complexes were eluted from the magnetic beads with 0.5 M NH₄OH (pH 11.0) containing 0.5 mM EDTA for 10 min under continuous mixing. Eluates were then submitted to the Proteomics and Lipidomics Core Facility of Weill Cornell Medicine for LC-MS/MS analysis following standard methods.

#### Animals

C57BL/6 WT male and female mice (#000664), Adipoq-Cre (#028020), BST2-floxed mice (B6.Cg-*Bst2tm1.1Bsz*/J, #029892), and hBST2 transgenic mice (B6.Cg-*Gt(ROSA)26Sortm1(CAG-BST2,-EGFP)Bsz*/J, #029893) were acquired from The Jackson Laboratory. Adipose tissue-specific BST2 knockout (Adipoq-Cre × BST2-floxed) and adipose tissue-specific hBST2 overexpressing (Adipoq-Cre × hBST2 transgenic) mice were generated by crossing the respective strains with Adipoq-Cre mice. All mice were maintained in a pathogen-free barrier-protected environment (12:12 h light/dark cycle, 22–24°C) at the Mount Sinai animal facilities. Mice were fed either a chow diet (Purina, 5001), a 60% kcal high-fat diet (HFD; Research Diets, D12492), or a high sucrose Western diet (WD; Research Diets, D12079B), with unrestricted access to food and water. Animal experiments were conducted in accordance with the Mount Sinai Institutional Animal Care and Research Advisory Committee. We did not use randomization in our allocation process since experimental groups were primarily separated based on genotype or sex, and investigators were not blinded to group allocation.

#### Glucose tolerance tests (GTT) and insulin tolerance tests (ITT)

For glucose tolerance tests (GTT), mice were fasted overnight and then injected intraperitoneally with glucose (1.5 g per kg body weight for males and 0.75 g per kg body weight for females). For insulin tolerance tests (ITT), mice were fasted for 6 h and then injected intraperitoneally with insulin (ReliOn/Novolin R, Novo Nordisk; 1.5 U per kg body weight). Blood samples were collected from a tail nick at the indicated time points, and glucose levels were measured using a glucometer with CONTOUR test strips (Ascensia Diabetes Care).

#### Retroviral overexpression of BST2 in 10T1/2 cells

cDNA encoding BST2, hBST2, and FLAG-BST2 was cloned into the pDONR221 entry vector using the Gateway Cloning System (Thermo Fisher Scientific). The sequences in the entry clones were then transferred by LR recombination into the pBabe-puro retroviral vector. pBabe plasmids were transfected into retrovirus packaging Phoenix-E cells for 48 h. Target cells (10T1/2) were plated at 50% confluency 24 h post-transfection. Forty-eight hours after transfection, media from transfected Phoenix-E cells were harvested and centrifuged for 5 min at 5,000 rpm to pellet cells and debris. Retrovirus-containing supernatant was carefully transferred onto target cells with polybrene (1:1000 dilution) and incubated overnight. The following day, media was replaced with regular growth media, and cells were incubated for an additional 24 h. Puromycin selection (4.5 μg/mL) was then performed to select for cells stably expressing BST2, hBST2, or FLAG-BST2.

### Lactate Assay In-Vitro

10T ½ cells, including BST2 Overexpression and Knock-out cells, were processed with Lactate Activity Assay (Sigma-Aldrich, MAK570). Cells were homogenized in 4 volumes of the provided Assay Buffer. LDH was deprotenized from the samples using 10 kDa MWCO spin filters (Thermo Scientific, 88513). Samples were prepared in 96-well plates at a 2 uL volume and brought to 50 uL/well with assay buffer. After incubating covered for 30 minutes at room temperature, absorbance was measured at 570 nm.

### G6P Activity Assay

10T ½ cells, including BST2 Overexpression and Knock-out cells, were processed with G6P Activity Assay (Sigma-Aldrich, MAK503) per manufacturer’s protocol. Samples were homogenized in 100 uL PBS and centrifuged at 14,000 RPM. 20 uL of each sample was placed in a 96-well plate, then brought to 100 uL using the working reagent. After incubating covered for 20 minutes at room temperature, the optical density was measured at 460 nm.

### SDD-AGE (Semi-Denaturing Detergent Agarose Gel Electrophoresis)

For SDD-AGE, crude mitochondria were suspended in sample buffer (0.5 x TBE containing 10% Glycerol, 2% SDS, and 0.0025% Bromophenol Blue). Mitochondria were isolated from 10T ½ cells using a kit for isolation of mitochondria from cultured cells per the manufacturer’s protocol (Thermo Fisher Scientific, 89874). Samples were loaded into a 1.5% agarose TBE gel containing 0.1% SDS in 1 x TBE running buffer containing 0.1% SDS. Electrophoresis was run vertically for 35 min at a constant 100V at 4 °C, then transferred to a Nitrocellulose Blotting Membrane (Amersham) for immunoblotting. For transfer, at 4 °C, a square of blotting membrane was cut to the size of the gel and soaked in 1x TBE. A stack of 20 pieces of mini-size Transfer Stacks (Bio-Rad, 12023835), 4 pieces of the Transfer Stack paper were soaked in 1xTBS. 1 piece of the soaked paper was added to the stack, followed by the Nitrocellulose membrane. The gel was rinsed in deionized water and added to the stack, followed by the remaining 3 pieces of soaked paper. Two trays filled with 1 x TBE were placed elevated beside the stack. A piece of Midi-size Transfer Stack paper (Bio-Rad, 12023956) pre-soaked in 1 x TBE was run from each of the elevated trays to the stack. A 500 mL bottle of water was placed atop the stack and allowed to proceed overnight. Staining was conducted identically to the Western Blot procedure.

### Extracellular Flux Analysis

Extracellular acidification rates (ECAR) during a modified Glycolysis Stress Test (Agilent #103020) were measured using a Seahorse XFe96 Extracellular Flux Analyzer (Agilent) according to the manufacturer’s instructions. Cells were seeded into XFe96 cell culture microplates (Agilent) at a density of 3,000 cells per well in 150 µL of complete DMEM supplemented with 10% FBS and allowed to fully differentiate into beige/white adipocytes prior to the assay. On the day of the assay, cells were washed three times and incubated in 180 µL of XF Base Medium (Agilent) supplemented with 2mM GlutaMax, adjusted to pH 7.4. Cells were equilibrated for 45 minutes at 37 °C in a non-CO₂ incubator prior to the assay. Extracellular acidification rates were measured basally and following sequential injections of norepinephrine (Port A, 2μM), glucose (Port B, 10 mM), oligomycin (Port C, 2 µM), and 2-deoxy-D-glucose (Port D, 50 mM). Glycolysis, glycolytic capacity, and non-glycolytic acidification were determined from extracellular acidification rate measurements, per manufacturer instructions.

### Lipidomics Analysis

Lipidomic profiling was performed by the UCLA Lipidomics Laboratory using differentiated 10T1/2 adipocytes. Cell pellets were collected after full adipocyte differentiation and submitted to the UCLA Lipidomics Core for targeted shotgun lipidomics analysis. Lipid extraction, sample preparation, and mass spectrometry analyses were performed according to the core facility’s standardized workflow. Quantitative lipid analysis was conducted using differential mobility spectrometry (DMS)-based shotgun lipidomics on a SCIEX 5500 triple quadrupole mass spectrometer equipped with a SelexION differential mobility device. Lipid species were analyzed using the Lipidyzer platform with internal lipid standards for quantitative measurements. This platform enabled the detection and quantification of multiple lipid classes, including triglycerides, phospholipids, sphingolipids, ceramides, cholesterol esters, and free fatty acids. Data acquisition and lipid quantification were performed using the Shotgun Lipidomics Assistant workflow as previously described. Lipid abundance was normalized as indicated in the corresponding figure legends.

### Quantification and statistical analysis

Data are presented as mean ± s.e.m. unless otherwise indicated. Each experiment was performed using independent biological replicates, and the number of biological replicates is indicated in the corresponding figure legends. No statistical method was used to predetermine sample size unless otherwise stated. Sample sizes were chosen based on prior experimental experience with comparable assays. Animals or samples were excluded only according to predefined technical criteria, including failed viral infection, failed RNA or protein quality control, or instrument malfunction. Statistical analyses were performed using GraphPad Prism. For comparisons between two groups, two-tailed unpaired Student’s t-tests were used unless paired analysis was appropriate. For comparisons among more than two groups, one-way or two-way analysis of variance was used followed by correction for multiple comparisons. Repeated-measures analyses were used for longitudinal measurements when appropriate. For indirect calorimetry, energy expenditure was analyzed using CalR where appropriate to account for differences in body mass or lean mass. P < 0.05 was considered statistically significant.

## Data availability

All sequencing and proteomics datasets generated in this study will be deposited in appropriate public repositories prior to publication. Source data for graphs and uncropped immunoblots will be provided with the final manuscript.

## Acknowledgements

Lipidomics was performed by Dr. Kevin Williams at the UCLA Lipidomics Core, microscopy was performed at ISMMS Microscopy Core, and Metabolomics was performed by ISMMS Metabolomics Core (RRID:SCR_027540) P.R is supported by R01DK136035, DP1DK140003, Irma T. Hirschl/Monique Weill-Caulier Foundation Scholar Award. L.G. is supported by AHA 25TPA1480256. C.H.C supported by F32DK141191. N.F.B supported by T32HL176457. The funders had no role in study design, data collection and interpretation, or the decision to submit the work for publication.

## Supplementary Figures

**Figure S1.**
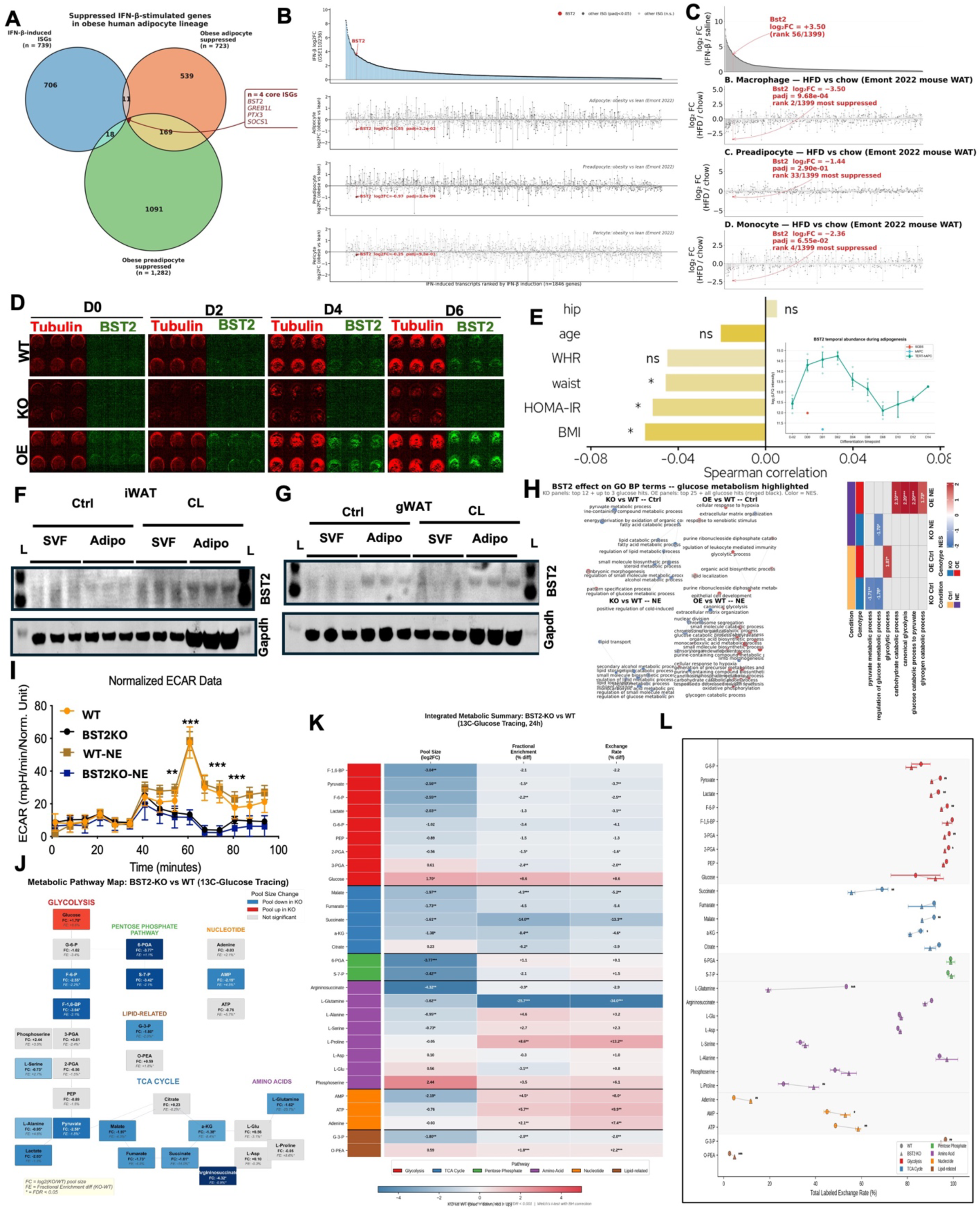
BST2 is selectively suppressed in obese adipocyte lineages and is required for adipocyte glycolytic and adipogenic chromatin programs. (A) Venn diagram of IFN-β-induced ISGs (n = 739), obese-adipocyte–suppressed genes (n = 723) and obese-preadipocyte–suppressed genes (n = 1,282); the four-gene core (BST2, GREB1L, PTX3, SOCS1) at the triple intersection is highlighted. (B) Ranked log₂FC of IFN-β-induced transcripts in human adipocyte, preadipocyte and pericyte lineages (Emont 2022 atlas, obese vs lean). BST2 is highlighted in red. (C) Ranked Bst2 log₂FC in macrophage, preadipocyte and monocyte fractions of mouse WAT (HFD vs chow), with adjusted p-values and rank in the IFN signature. (D) Time-course in cell western blot (D0, D2, D4, D6) for Tubulin and BST2 in WT, BST2KO and BST2OE cells; replicates per condition shown. (E) Spearman correlation of BST2 expression with clinical metabolic parameters (BMI, HOMA-IR, waist, WHR, age, hip) in a human-adipose cohort (* FDR < 0.05). Inset: BST2 protein abundance time-course during adipogenesis in iWAT-PreAd, MAPC and TERT-MAPC lines. (F) Western blot of BST2 and GAPDH in SVF and mature-adipocyte fractions from iWAT under Ctrl vs CL-316,243 treatment. (G) Same as (F) but for gWAT. (H) Gene Ontology BP enrichment word clouds and clustered heatmap for KO-vs-WT and OE-vs-WT comparisons under Ctrl and NE; glucose-metabolism terms are highlighted (NES with significance asterisks). (I) Replicate Seahorse ECAR glycolysis-stress test in WT, BST2KO, WT-NE and BST2KO-NE adipocytes. **p < 0.01, ***p < 0.001 at indicated time-points (mean ± s.e.m., n= 5–6 wells). (J) Metabolic pathway map (glycolysis, PPP, TCA, nucleotide, amino-acid, lipid-related) annotated with pool-size log₂FC (KO/WT) and fractional-enrichment differences from ¹³C-glucose tracing (24 h). (K) Integrated metabolic summary heatmap of pool size (log₂FC), fractional enrichment (% diff) and exchange rate (% diff) in BST2KO vs WT across ∼30 metabolites, grouped by pathway. FDR < 0.05, FDR < 0.01, ** FDR < 0.001 (Welch t-test with BH correction). (L) Scatter plot of total labelled exchange rate (%) for each metabolite (mean ± s.e.m.) in WT (circles) vs BST2KO (triangles), colour-coded by pathway.

**Figure S2.**
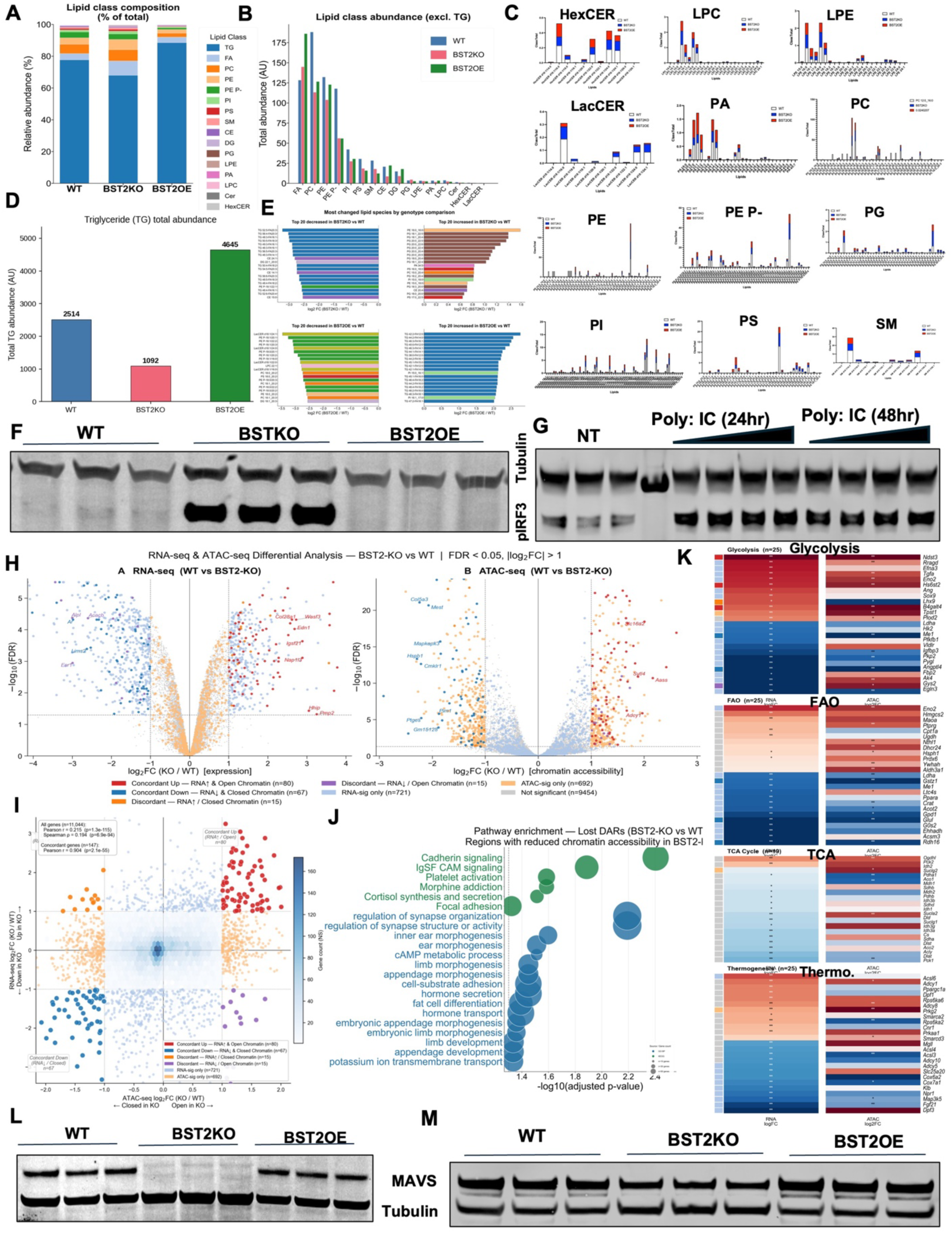
BST2 selectively controls lipid and innate immune signaling. (A) Stacked bar chart of lipid-class composition (% of total) in WT, BST2KO and BST2OE. (B) Lipid-class abundance (excluding TG) across the three genotypes. (C) Species-level violin/box plots for HexCER, LPC, LPE, LacCER, PA, PC, PE, PE P-, PG, PI, PS and SM. (D) Total triglyceride abundance bar chart across genotypes. (E) Top-20 most up- and down-regulated lipid species in BST2KO vs WT and BST2OE vs WT (log₂FC). (F) Western blot of Tubulin and pIRF3 in WT and BST2OE adipocytes. (G) Western blot of Tubulin and pIRF3 in WT and BST2OE adipocytes under NT, poly(I:C) 24 h and poly(I:C) 48 h. (H) Paired volcano plots of RNA-seq and ATAC-seq differential analyses for BST2KO vs WT (FDR < 0.05, |log₂FC| > 1) with concordance color-coding. (I) Scatter of ATAC-seq log₂FC vs RNA-seq log₂FC for all genes with Pearson/Spearman statistics. (J) Pathway-enrichment bubble plot of regions losing accessibility in BST2KO (fat-cell differentiation, cadherin/IgSF CAM signalling, morphogenesis). (K) Concordance heatmap of RNA-seq and ATAC-seq log₂FC for genes in Glycolysis, FAO, TCA Cycle and Thermogenesis hallmark sets, with gene labels. (L and M) Western blot of MAVS and Tubulin in WT, BST2KO and BST2OE differentiated adipocytes (L) and preadipocytes (M).

**Figure S3.**
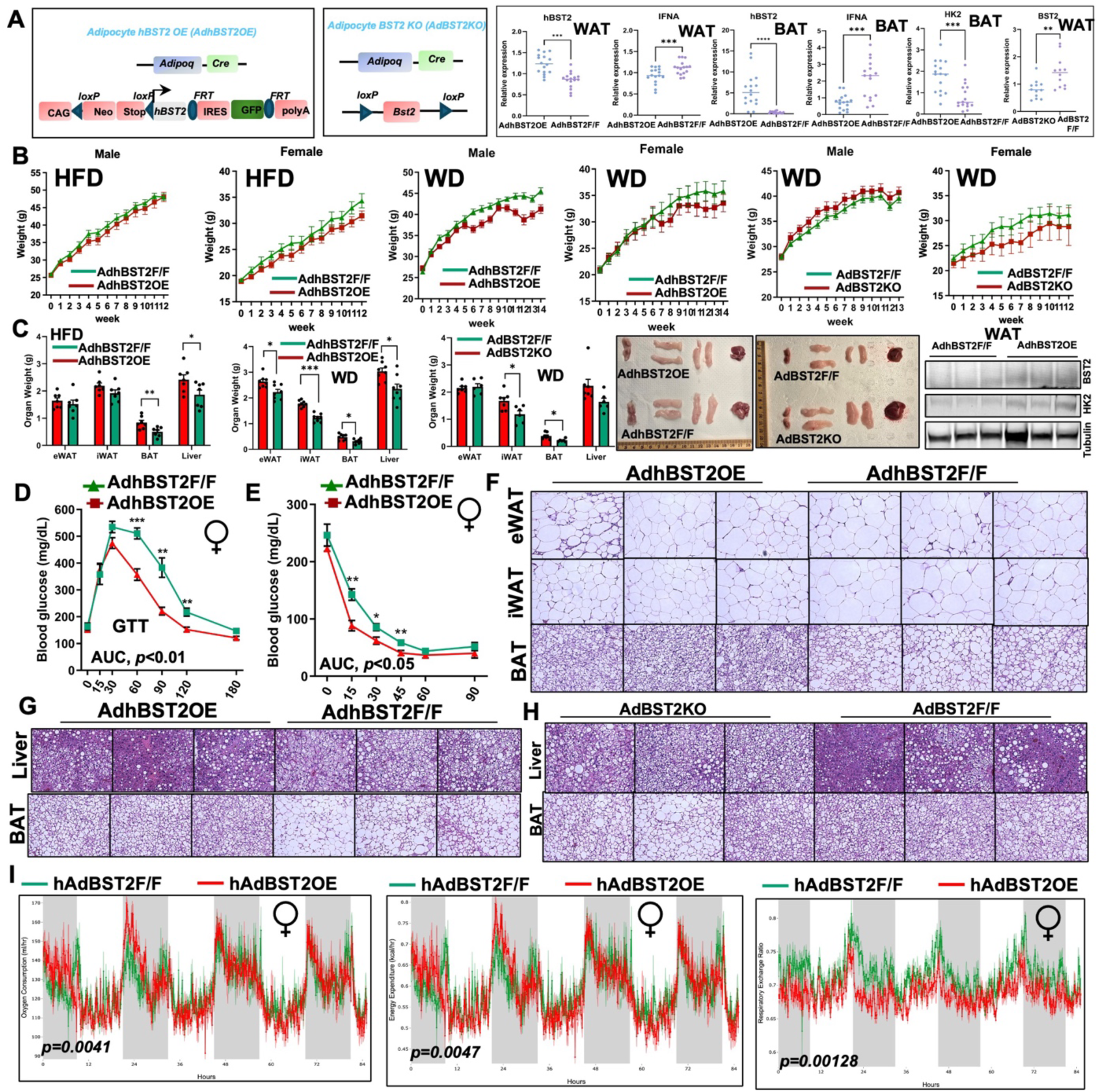
Generation and extended in vivo characterization of adipocyte-specific BST2 gain- and loss-of-function models. (A) Genetic strategy schematics for AdhBST2OE (CAG-loxP-Neo-Stop-loxP-hBST2-IRES-GFP-FRT-polyA × Adipoq-Cre) and AdBST2KO (Bst2ᶠ/ᶠ × Adipoq-Cre). Real time qPCR for indicated genes in WAT of AdhBST2OE or AdBST2KO mice. (B) Body-weight curves over 10–14 weeks under HFD (Males and Females) and high sucrose Western diet (WD) (Males and Females) for AdhBST2OE vs AdhBST2ᶠ/ᶠ and AdBST2KO vs AdBST2ᶠ/ᶠ (6 sub-panels). (C) Organ-weight bar charts, gross pathology (eWAT, iWAT, BAT, liver), and western blot in indicated mouse lines. (D) GTT in 12 Weeks HFD-fed female AdhBST2OE vs AdhBST2ᶠ/ᶠ; AUC p < 0.01. (E) ITT in 12 Weeks HFD-fed female AdhBST2OE vs AdhBST2ᶠ/ᶠ; AUC p < 0.05. (F) H&E histology for BAT, iWAT, and eWAT of 12 Weeks HFD-fed AdhBST2OE vs AdhBST2ᶠ/ᶠ. (G) H&E histology of liver and BAT from 12 Weeks WD-fed AdhBST2OE vs AdhBST2ᶠ/ᶠ. (H) H&E histology of liver and BAT from 12 Weeks WD-fed AdBST2KO vs AdBST2ᶠ/ᶠ. (I) Female CLAMS time-series over 72 h (light/dark shading) for hAdBST2OE vs hAdBST2ᶠ/ᶠ: VO₂ (p = 0.0041), energy expenditure (p = 0.0047) and RER (p = 0.00128).

**Figure S4.**
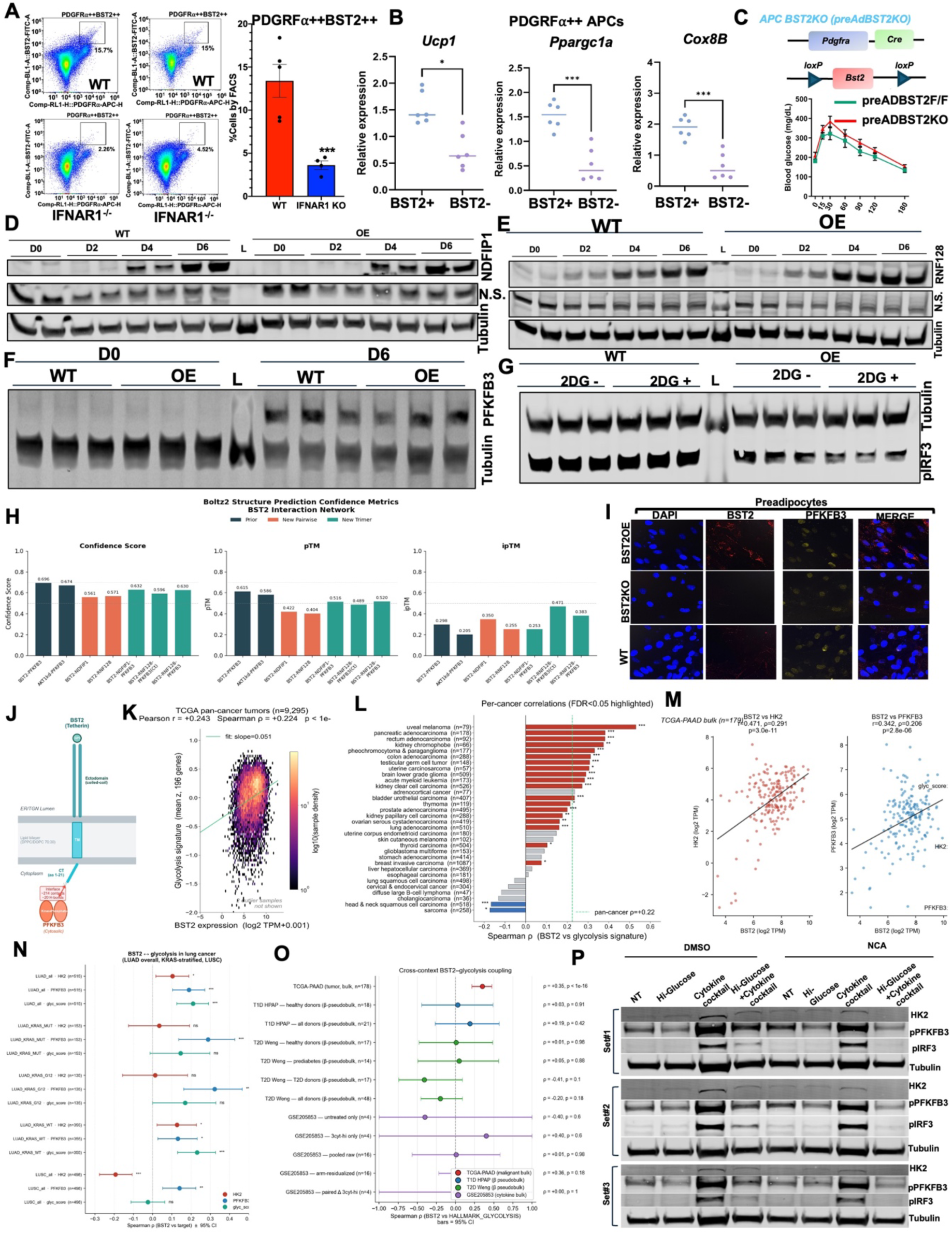
Mechanistic, structural and translational validation of the BST2 immunometabolic checkpoint. (A) Flow cytometry of PDGFRα⁺⁺ BST2⁺⁺ cells in WT and Ifnar1⁻/⁻ adipose with quantification. BST2⁺⁺ frequency falls from 13.6% in WT to 3.4% in Ifnar1⁻/⁻ (*** p< 0.001). (B) qPCR of Ucp1, Ppargc1a and Cox8b in PDGFRα⁺BST2+ or PDGFRα⁺BST2- APCs sorted by BST2 status. * p < 0.05, ***p < 0.001 (BST2⁺ vs BST2⁻). (C) Schematic of preAdBST2KO model (Pdgfra-Cre × Bst2ᶠ/ᶠ) and GTT showing glucose tolerance vs littermate controls. (D) Time-course Western blot of NDFIP1 and Tubulin (WT vs OE; D0, D2, D4, D6) in WT and BST2OE adipocytes; a non-specific band (N.S.) is indicated. (E) Time-course Western blot of RNF128 and Tubulin in WT and BST2OE adipocytes (D0–D6); a non-specific band (N.S.) is indicated. (F) Time-course Western blot of PFKFB3 and Tubulin (WT vs OE; D0 and D6). (G) Western blot of pIRF3 and Tubulin in WT and BST2OE adipocytes ± 2-deoxyglucose (2-DG, 10 mM). (H) Boltz2 structure-prediction confidence metrics (Confidence Score, pTM, ipTM) for BST2 binary and trimeric complexes with PFKFB3, NDFIP1, RNF128 and combinations. (I) IF of BST2OE, BST2KO and WT preadipocytes stained for DAPI (blue), BST2 (red), PFKFB3 (yellow) and MERGE. (J) Schematic of BST2 (Tetherin) membrane topology and its interaction network with PFKFB3. (K) TCGA pan-cancer scatter (n = 9,295) of BST2 expression vs Hallmark Glycolysis signature z-score. Pearson r = 0.243; Spearman ρ = 0.224; p < 1 × 10⁻. (L) Forest plot of per-cancer Spearman ρ between BST2 and glycolysis signature across ∼33 TCGA tumor types (FDR < 0.05 highlighted, red bars). Pan-cancer ρ = 0.22. (M) TCGA-PAAD (n = 178) bulk correlation scatters: BST2 vs HK2 (r = 0.471, p = 3.0 × 10⁻¹¹) and BST2 vs PFKFB3 (r = 0.342, p = 2.6 × 10⁻⁶). (N) Forest plot of BST2–glycolysis correlations (HK2, PFKFB3, Hallmark Glycolysis score) in LUAD (all, KRAS-MUT, KRAS-WT) and LUSC. (O) Cross-context BST2–glycolysis Spearman ρ with 95% CI across TCGA-PAAD bulk, T1D/T2D pancreatic pseudobulk (HPAP, Weng) and GEO adipocyte datasets (GSE205853). (P) Western blot of HK2, pPFKFB3, pIRF3 and Tubulin in three biological replicate sets under NT, Hi-Glucose, cytokine cocktail and Hi-Glucose + cytokine, with DMSO or 10μM NCA (BST2 inhibitor) treatment.

