## Supplementary Tables for "Tetherin enforces an immunometabolic checkpoint that coordinates glycolytic and interferon signaling in adipocytes"

**Supplementary Table 1. Real-time PCR Primers**

| **PRIMER** | **FORWARD** | **REVERSE** |
| --- | --- | --- |
| Gapdh | CATCACTGCCACCCAGAAGACTG | ATGCCAGTGAGCTTCCCGTTCAG |
| Bst2 (mouse) | CAAACTCCTGCAACCTGACCGT | CTCCTGGTTCAGCTTCGTGACT |
| BST2 (human) | TCTCCTGCAACAAGAGCTGACC | TCTCTGCATCCAGGGAAGCCAT |
| Hk2 | CCCTGTGAAGATGTTGCCCACT | CCTTCGCTTGCCATTACGCACG |
| Pparg | AACTCTGGGAGATTCTCCTGTTGA | TGGTAATTTCTTGTGAAGTGCTCATA |
| Ppara | GACAAGGCCTCAGGGTACCA | GCCGAATAGTTCGCCGAAA |
| Ucp1 | GGCCTCTACGACTCAGTCCA | TAAGCCGGCTGAGATCTTGT |
| Ifna1 | GGATGTGACCTTCCTCAGACTC | ACCTTCTCCTGCGGGAATCCAA |
| Adipoq | CCGGAACCCCTGGCAG | CTGAACGCTGAGCGATACACA |

**Supplementary Table 2. Antibodies**

| Reagent | Source | Catalogue No. | Working Conc. |
| --- | --- | --- | --- |
| BST2 | Novus Biologicals | DDX0390P-100 | IF |
| IRF-3 | Cell Signaling | 4302S | WB (1:1000) |
| Phospho-IRF-3 (Ser396) | Cell Signaling | 4947S | WB (1:1000) |
| Hexokinase II | Cell Signaling | 2867S | WB (1:1000) |
| PFKFB3 | Cell Signaling | 13123T | WB (1:1000) |
| Anti-PFKFB3 | Abcam | ab181861 | IF (1:500) |
| Phospho-PFKFB3 (Ser461) | Invitrogen | PA5-114619 | WB (1:1000)  IF (1:500) |
| STING | Cell Signaling | 13647T | WB (1:1000) |
| Phospho-STING (Ser365) | Cell Signaling | 72971T | WB (1:1000) |
| TBK1/NAK | Cell Signaling | 3504T | WB (1:1000) |
| Phospho-TBK1/NAK (Ser172) | Cell Signaling | 5483S | WB (1:1000) |
| RNF128 | Proteintech | 26015-1-AP | WB (1:1000) |
| NDFIP1 | Proteintech | 15602-1-AP | WB (1:1000) |
| Anti-α-Tubulin | MilliporeSigma | CP06-100UG | WB (1:1000) |
| IRDye® 800CW Goat anti-Rabbit IgG Secondary Antibody | LICORBio | 926-32211 | WB (1:10000) |
| IRDye® 680RD Goat anti-Mouse IgG Secondary Antibody | LICORBio | 926-68070 | WB (1:10000) |
| Goat Anti-Rat IgG H&L (Alexa Fluor® 647) | Abcam | ab150159 | IF (1:500) |
| Goat Anti-Rabbit IgG H&L (Alexa Fluor® 568) | Abcam | ab175471 | IF (1:500) |

**Supplementary Table 3. sgRNA Sequences Used for CRISPR–Cas9 Genome Editing**

| **sgRNA** | **Sequence (5' to 3')** |
| --- | --- |
| Ndfip1 #1 | AACACTGCCCAGTTATGACG |
| Ndfip1 #2 | GCTACGATCCCTTGTTCC |
| Ndfip1 #3 | ACATTATACGATGGGGCTT |
| Ndfip2 #1 | TAACTCAGAATCATCTGCTG |
| Ndfip2 #2 | GCTGCTAAGCTAGAAGTTGA |
| Rnf128 #1 | CACGGTGCAAGTATCTTGGT |
| Rnf128 #2 | ATTTCACGGTGCCCACGGTT |
| Rnf128 #3 | AACCGTGGGCACCGTGAAAT |
| Pfkfb3 #1 | TGTAAGTCTTACCCCGGGCT |
| Pfkfb3 #2 | CACAATCACGGTTGGGGAGT |
| Bst2 #1 | TGACGGCGAAGTAGATTGTC |
| Bst2 #2 | GAACAGGACCACCAAGATCG |
| Bst2 #3 | TGAAGTCACGAAGCTGAACC |
